# Genomics of sorghum local adaptation to a parasitic plant

**DOI:** 10.1101/633529

**Authors:** Emily S. Bellis, Elizabeth A. Kelly, Claire M. Lorts, Huirong Gao, Victoria L. DeLeo, Germinal Rouhan, Andrew Budden, Govinal Badiger Bhaskara, Zhenbin Hu, Robert Muscarella, Michael P. Timko, Baloua Nebie, Steven M. Runo, N. Doane Chilcoat, Thomas E. Juenger, Geoffrey P. Morris, Claude W. dePamphilis, Jesse R. Lasky

## Abstract

Host-parasite coevolution can maintain high levels of genetic diversity in traits involved in species interactions. In many systems, host traits exploited by parasites are constrained by use in other functions, leading to complex selective pressures across space and time. Here, we study genome-wide variation in the staple crop *Sorghum bicolor* (L.) Moench and its association with the parasitic weed *Striga hermonthica* (Delile) Benth., a major constraint to food security in Africa. We hypothesize that geographic selection mosaics across gradients of parasite occurrence maintain genetic diversity in sorghum landrace resistance. Suggesting a role in local adaptation to parasite pressure, multiple independent loss-of-function alleles at sorghum *LOW GERMINATION STIMULANT 1 (LGS1)* are broadly distributed among African landraces and geographically associated with *S. hermonthica* occurrence. However, low frequency of these alleles within *S. hermonthica*-prone regions and their absence elsewhere implicate potential tradeoffs restricting their fixation. *LGS1* is thought to cause resistance by changing stereochemistry of strigolactones, hormones that control plant architecture and belowground signaling to mycorrhizae and are required to stimulate parasite germination. Consistent with tradeoffs, we find signatures of balancing selection surrounding *LGS1* and other candidates from analysis of genome-wide associations with parasite distribution. Experiments with CRISPR-Cas9 edited sorghum further indicate the benefit of *LGS1*-mediated resistance strongly depends on parasite genotype and abiotic environment and comes at the cost of reduced photosystem gene expression. Our study demonstrates long-term maintenance of diversity in host resistance genes across smallholder agroecosystems, providing a valuable comparison to both industrial farming systems and natural communities.

**SIGNIFICANCE STATEMENT:** Understanding co-evolution in crop-parasite systems is critical to management of myriad pests and pathogens confronting modern agriculture. In contrast to wild plant communities, parasites in agricultural ecosystems are usually expected to gain the upper hand in co-evolutionary ‘arms races’ due to limited genetic diversity of host crops in cultivation. Here, we develop a framework to characterize associations between genome variants in global landraces (traditional varieties) of the staple crop sorghum with the distribution of the devastating parasitic weed *Striga hermonthica.* We find long-term maintenance of diversity in genes related to parasite resistance, highlighting an important role of host adaptation for co-evolutionary dynamics in smallholder agroecosystems.

## INTRODUCTION

Host-parasite interactions can be powerful and dynamic selective forces maintaining genetic variation in natural populations (1). In wild plant pathosystems, long-term balancing selection often maintains diverse resistance alleles in host populations (2–4). When rare alleles provide a selective advantage, negative frequency-dependent selection drives cycling of resistance and virulence alleles (i.e. fluctuating Red Queen dynamics *sensu* (5)) (2, 6). Fitness costs of resistance and spatiotemporal changes in selection can also maintain diversity across gradients of parasite pressure (7).

In contrast to wild plant communities where fluctuating Red Queen dynamics have frequently been observed, low host diversity in agricultural settings is often assumed to permit runaway ‘arms races’ in fast-evolving parasites (4, 8). Relative to smallholder farms, however, industrial scale farming accounts for a fraction of global production for many food crops (9). The dynamics of host adaptation to parasites across diverse smallholder agricultural systems remains poorly known, despite relevance for identifying novel resistance alleles and managing crop genetic resources (e.g. preserving germplasm both *ex situ* and *in situ*; (10)). Are co-evolutionary dynamics in smallholder farming systems more similar to natural plant pathosystems, where high connectivity among genetically diverse patches can help promote evolution of host resistance (11)?

Current approaches for identifying and studying the evolution of resistance alleles often involve scoring large panels of diverse individuals (in genome-wide association studies, GWAS) or many recombinant individuals deriving from controlled crosses (in linkage mapping) and using DNA sequence information to identify genomic regions associated with parasite susceptibility. These mapping studies have revealed numerous insights to mechanisms of plant-pathogen dynamics (12), but require extensive phenotyping and genotyping effort for adequate statistical power. By contrast, if traditional local crop or livestock varieties (known as landraces) are locally adapted to regions of high parasite prevalence (13–16), then a different ‘bottom-up’ approach may be used. Compared to modern improved varieties, which may have lost resistance alleles due to bottlenecks and selection in optimal environments (17), landraces and wild relatives may be a rich source of resistance alleles to sympatric parasites. Specifically, to map loci underlying local adaptation, one can identify loci where allele frequency is associated with environmental conditions, known as genotype-environment associations (GEA). GEA of georeferenced landraces have been a powerful strategy for understanding the genetic basis of local adaptation to gradients of abiotic stressors (14, 15). Furthermore, landraces can be studied to test the hypothesis that a putatively environmentally-adapted allele, identified from a limited set of experimental environments and genetic backgrounds, is indeed adaptive across a wide range of similar environments and across diverse genetic backgrounds (14).

In this study, we extend GEA to biotic stress gradients to evaluate the frequency and distribution of alleles in genomes of sorghum that confer resistance to the African witchweed *Striga hermonthica* (Delile) Benth., a root hemiparasite of the broomrape family (Orobanchaceae)*. Sorghum bicolor* (L.) Moench is the world’s fifth most important crop and was domesticated from the wild progenitor *Sorghum arundinaceum* (Desv.) Stapf in Africa more than 5000 years ago (18). Sorghum is particularly important due to its tolerance of marginal environments compared to maize and rice. While plant responses to environment are influenced by many processes, in recent years the role of strigolactones, hormones that regulate shoot branching (19), root architecture (20) and response to abiotic stress (21), have received increasing attention. Strigolactones exuded from roots, particularly under nutrient (22) and water (23) limitation, promote interactions with beneficial arbuscular mycorrhizal fungi (24) and also function as germination stimulants for many root-parasitic plants. *S. hermonthica* attacks cereal crops and is one of the greatest biotic threats to food security in Africa, causing $US billions in crop losses annually (25). Several resistance mechanisms have been reported among cultivated and wild sorghums (26).

Here, we evaluate the hypothesis that a geographical selection mosaic across gradients of biotic interactions maintains genetic diversity in sorghum landrace resistance to *S. hermonthica* (27, 28). To identify genomic signatures of host adaptation to parasites, we first develop species distribution models for *S. hermonthica* and search for statistical associations between sorghum genotype and modeled parasite prevalence at the location of origin for each sorghum landrace. We validate our approach by characterizing diversity and geographic distribution of loss-of-function alleles at the putative sorghum resistance gene *LOW GERMINATION STIMULANT 1* (*LGS1*) (29). Loss-of-function mutations at *LGS1*, a gene of unknown function with a sulfotransferase domain, is thought to underlie a quantitative trait locus (QTL) that alters stereochemistry of the dominant strigolactone in sorghum root exudates, from 5-deoxystrigol to orobanchol (29). Orobanchol is considered a weaker stimulant of *S. hermonthica* germination, conferring resistance (29, 30). Our ecological genetic analyses of diverse sorghum landraces suggest that *LGS1* loss-of-function mutations are adaptive across a large region of high *S. hermonthica* prevalence in Africa. However, using comparisons of CRISPR-Cas9 edited sorghum, we also present evidence supporting potential trade-offs for *LGS1* loss-of-function due to high sensitivity of some *S. hermonthica* genotypes to orobanchol and subtle impacts on host fitness. In addition to focused analyses on *LGS1*, we perform genome-wide tests of association with parasite distribution. We investigate patterns of polymorphism surrounding candidate resistance genes with evidence of locally-adaptive natural variation to determine whether balancing selection has maintained diversity in *S. hermonthica* resistance over evolutionary time.

## RESULTS

### S. hermonthica *distribution model*

We predicted that host alleles conferring resistance or tolerance would be strongly associated with the geographic distribution of *Striga* parasites. To identify regions of likely *S. hermonthica* occurrence in the absence of continent-wide surveys, we built MaxEnt species distribution models (SDMs) (31). This approach uses locations of known species occurrences and multivariate environmental data to generate predictions of suitable habitat across a landscape. The optimal model showed good ability to predict occurrence with an AUC value of 0.86 (Fig. 1). A high degree of overlap was observed between models generated using all *S. hermonthica* records (*n*=1,050) and a subset of 262 occurrences that were observed specifically in fields of sorghum (Schoener’s *D* = 0.82; *I* = 0.97). Annual rainfall and total soil N were the most informative variables for predicting *S. hermonthica* occurrence (Table S1). Compared to all grid cells in the study background, distributions of environmental values for locations with high habitat suitability (HS) were generally restricted to a narrower range of intermediate values of precipitation and soil quality (Fig. S1-2). Locations with the highest HS scores exhibited mean annual rainfall ranging from ~500-1300 mm/year and soil nitrogen ranging from ~400 – 1000 g/kg (10^th^-90^th^ percentiles for all cells with HS >0.5, Table S1). Soil clay content also contributed substantially to the sorghum-only model, and clay content in locations with the highest HS scores ranged from 12-29% (10^th^-90^th^ percentile, HS > 0.5, all-occurrence model) or up to 36% (sorghum-only model, Table S1).

**Figure 1:**
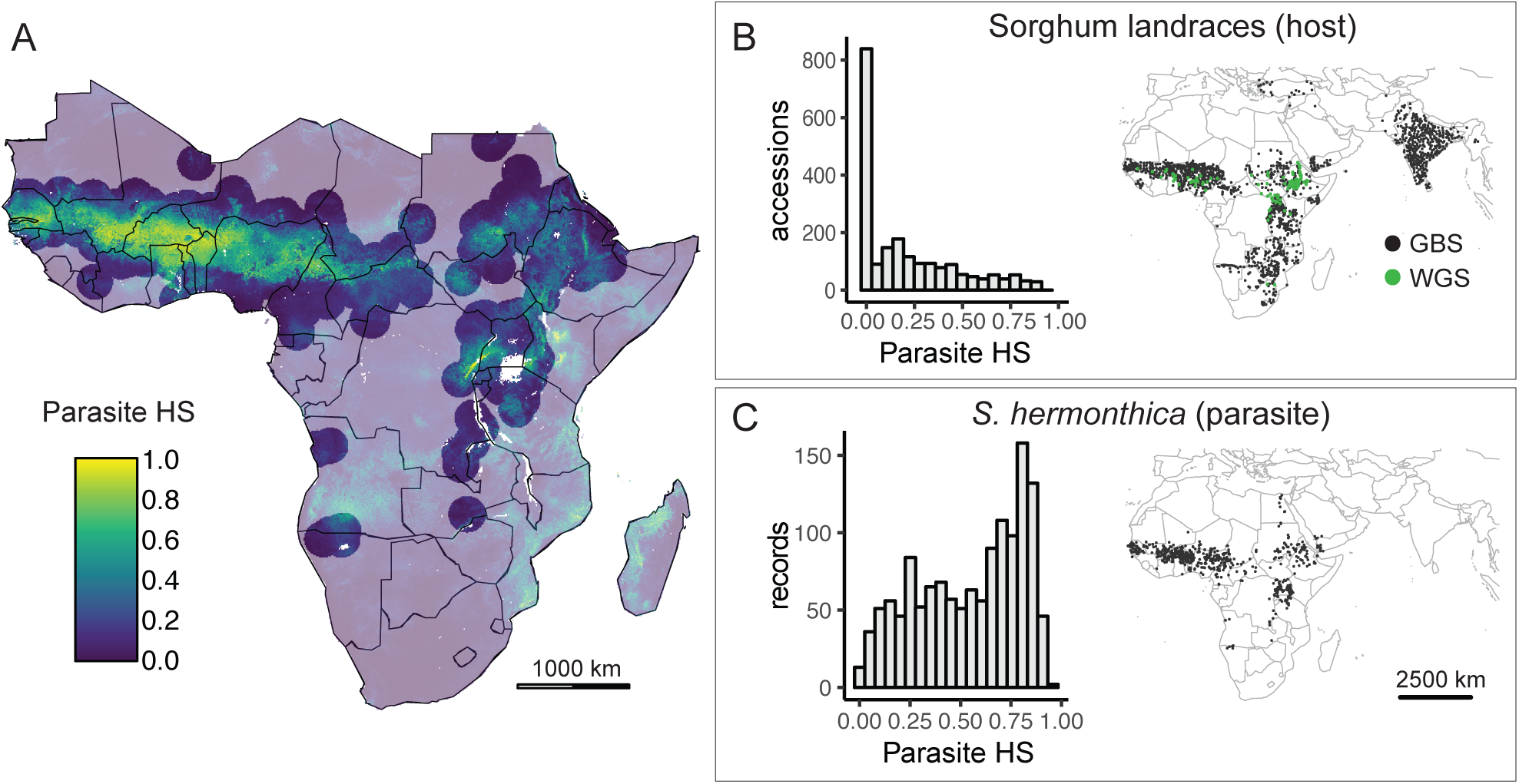
A subset of global sorghum landraces are from parasite-prone areas. A) *Striga hermonthica* habitat suitability (HS) scores across Africa based on MaxEnt species distribution model. To account for areas with suitable habitat where parasites have not been recorded, HS for accessions more than 200 km from any *S. hermonthica* record was set to zero; transparent colors demarcate areas of this HS masking. B) Geographic distribution and frequency histogram of HS scores at locations of all georeferenced and genotyped sorghum landrace accessions. Landraces with available genotyping-by-sequencing (GBS, *n=*2070) and whole genome sequence (WGS, *n=*143) data are shown. C) Geographic distribution and frequency histogram of HS scores at locations of 1369 *S. hermonthica* occurrence records.

### LGS1 *associations with* S. hermonthica *occurrence*

We predicted that sorghum resistance alleles would be more common in parasite-prone regions. Evaluating this prediction also allowed us to validate our species distribution model-based genotype-environment association approach by characterizing associations between *S. hermonthica* distribution and genetic variation at *LGS1* (*Sobic*.*005G213600*). *LGS1* is thought to cause a known QTL for resistance to *Striga* spp. (29).

Using whole genome sequencing data (~25x coverage) from 143 sorghum landraces, we found evidence for three naturally occurring mutations resulting in *LGS1* loss-of-function (Fig. 2). Two ~30 kb deletions were identified between positions 69,958,377-69,986,892 (*n* = 5 accessions) and 69,981,502-70,011,149 (*n* = 4 accessions, Table S2; Fig. 2A). These deletions appear to be identical to two previously described as resistance alleles, *lgs1-2* and *lgs1-3* (29), although breakpoint positions reported by our structural variant caller differed slightly. No SNPs in a separate genotyping-by-sequencing (GBS) dataset of >2000 sorghum landraces tagged *lgs1-2* or *lgs1-3* (Fig. S3), and so we imputed large deletions identified from the WGS dataset to the GBS dataset based on patterns of missing data (see Methods). Deletion imputations were validated by testing root exudate from a subset of sorghum accessions for their ability to induce *S. hermonthica* germination (Fig. S4). We tested four genotypes with likely deletion alleles, and these genotypes stimulated significantly fewer *Striga* seeds to germinate compared to eight other genotypes that did not show strong evidence for *lgs1-2* or *lgs1-3* deletions (linear mixed-effects model with genotype random effect, deletion genotypes stimulated germination of 8.43 fewer seeds out of 75 per replicate, Wald 95% CI=1.49,15.36).

**Figure 2.**
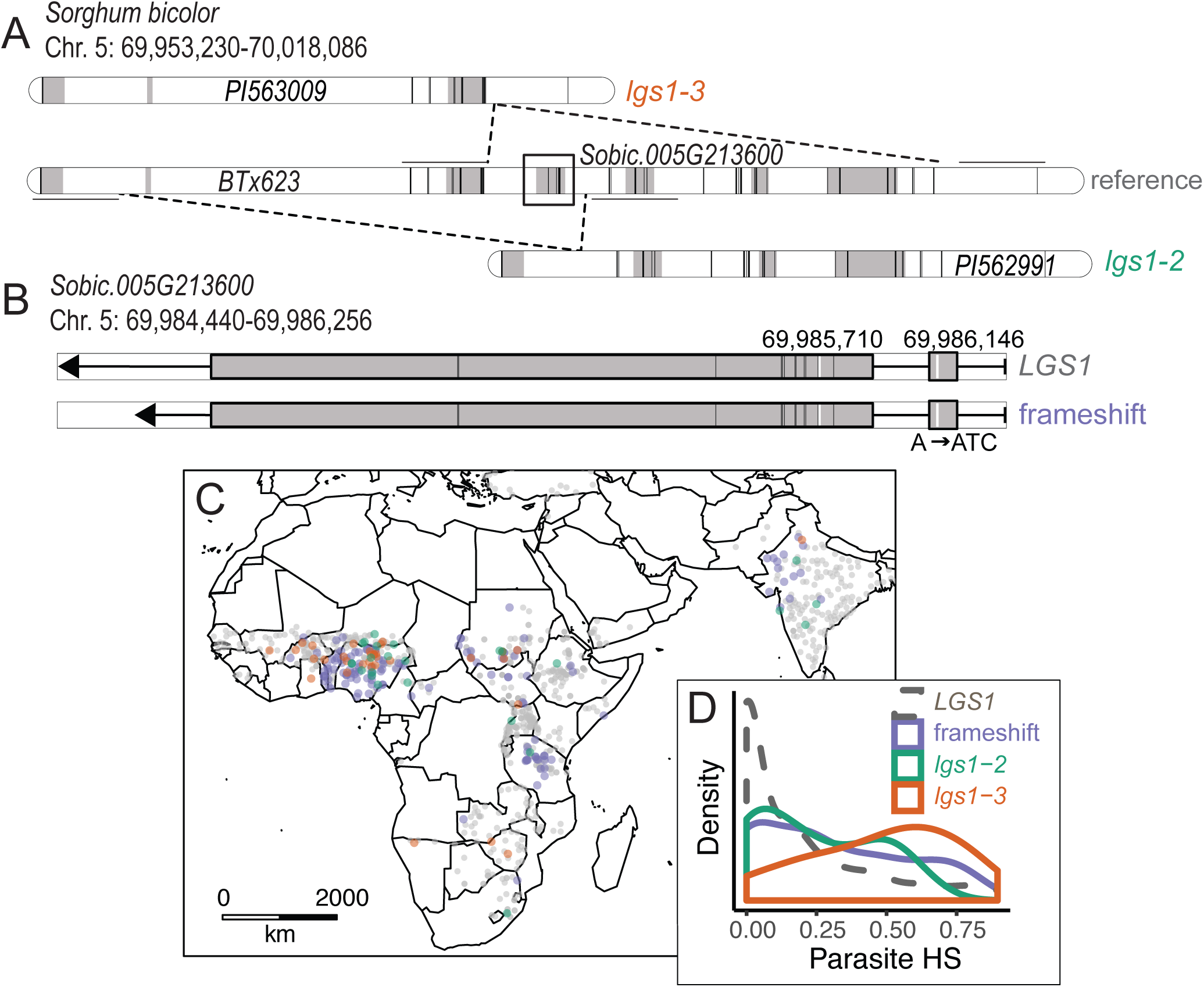
*LGS1* loss-of-function alleles are broadly distributed within parasite-prone regions. Schematic of large deletion variants (A) and frameshift mutation (B) impacting sorghum *LGS1*, a locus involved in resistance to *Striga hermonthica*. Grey shading indicates position of gene models (A) or coding regions (B). Vertical black bars indicate position of SNPs in the GBS dataset, and horizontal black lines denote 5 kb flanking regions used to impute deletion calls. In B, vertical white bars show the frameshift mutation (position 69,986,146) and the SNP at position 69,985,710 that tags the frameshift in the GBS dataset. C) Geographic distribution of *LGS1* alleles in sorghum landraces. D) Distribution of parasite habitat suitability (HS) scores at locations of sorghum accessions with *lgs1-2* (*n*=25), *lgs1-3* (*n*=34), frameshift (*n*=131), or intact *LGS1* (*n=*785).

In addition to *lgs1-2* and *lgs1-3*, we identified a previously unknown two bp insertion predicted to cause a frameshift variant in the beginning of the *LGS1* coding region (position 69986146, allele frequency in WGS dataset = 8%). The frameshift was linked to a SNP genotyped in the GBS dataset (Fig. 2B, T/A at position 69,985,710; D’ = 0.93, *r*^*2*^ = 0.84). All nine accessions with the frameshift in the WGS dataset also shared a 315 bp deletion (positions 69,984,268-69,984,583) overlapping 143 bp of the 3’-untranslated region in the 1580 bp second exon of *LGS1*.

Three of the six total independent *LGS1* putative loss-of-function alleles characterized here and elsewhere (29) were present at low frequency in the GBS panel. Among accessions with SNP calls in *LGS1*, 7.0% of accessions exhibited SNP calls consistent with homozygous *lgs1-2* and *lgs1-3* in the GBS dataset and the SNP tagging the frameshift was present at an allele frequency of 15%. (Supplemental Data File S1). *LGS1* loss-of-function alleles were found in diverse genetic backgrounds and geographic regions (Fig. S5, Table S2, Supplemental Data File S1), suggesting that these mutations have had time to spread, their benefit is not strongly masked by epistasis, and costs of resistance are not strong enough to prevent their spread. Most landraces with *LGS1* loss-of-function alleles and known botanical race (which largely correspond to genetic clusters (3)) from the GBS dataset were guinea (211 accessions) or caudatum (165 accessions, Supplemental Data File S1; 298 lacked assignments).

*LGS1* loss-of-function alleles were more common among landraces with high parasite HS scores (Fig. 2C-D). However, correlations with population structure reduced power to detect associations with these resistance alleles after accounting for kinship (Fig. S5). The median *S. hermonthica* HS score was 0.20 for accessions homozygous for *lgs1-2*, 0.54 for accessions homozygous for *lgs1-3*, 0.25 for accession with the frameshift, and 0.09 for accessions without evidence for *LGS1* loss-of-function. The difference in *S. hermonthica* HS score between *lgs1-3* and *LGS1* intact accessions was statistically significant before (*p* < 0.001, Wilcoxon rank sum test) but not after accounting for relatedness (*p =* 0.10, MLM). Frameshift associations with HS were also statistically significant prior to correction for relatedness (*p* < 0.001, Wilcoxon rank sum; *p =* 0.69, MLM). We observed modest support for associations between *lgs1-2* and *S. hermonthica* HS before correcting for relatedness (*p =* 0.06, Wilcoxon rank sum test; *p* = 0.53, MLM). Results were similar considering a subset of just African accessions (*p_lgs1-3_<0.001*, *p_frameshift_ <0.001*, and *p_lgs1-2_* = 0.07; Wilcoxon rank sum test). For sorghum landraces within parasite-prone regions (200 km from an occurrence record), naturally occurring *LGS1* loss-of-function alleles were more common among landraces from low nutrient (particularly low nitrogen) environments (*p_N_ < 0.001, p_P_* = 0.004; Kolmogorov-Smirnov test comparing distribution of soil N or P for *LGS1* intact vs. *LGS1* loss-of-function landraces) but evenly distributed across precipitation gradients (*p* = 0.4; Kolmogorov-Smirnov test).

### Parasite-associated SNPs across the sorghum genome

We performed genome-wide tests of association with predicted parasite HS score. A scan of 317,294 SNPs across 2070 sorghum landraces revealed 97 genomic regions exhibiting significant associations with *S. hermonthica* distribution at a false discovery rate of 5% (Fig. 3A, Table S4). Of SNPs exceeding the threshold for significance, 45 were present within 1 kb of a predicted gene model (Table S4). Three SNPs exceeding this threshold were in QTL previously associated with *Striga* resistance (32) including one intron variant in the uncharacterized gene model *Sobic.001G227800* (Table S4). Another SNP was present in an intron of *Sobic.007G090900*, a gene model with high homology to *SMAX1/D53*, which is degraded in a strigolactone-dependent manner to control downstream SL signaling and is associated with tillering and height in rice (33). SNPs among those with the strongest associations to parasite occurrence were also found in genes related to suberin and wax ester biosynthesis (*Sobic.007G091200*, synonymous variant) including phenylalanine ammonia-lyase (*Sobic.006G148800*, intron variant). Phenylalanine ammonia-lyase is highly upregulated in the resistant rice line Nipponbare compared to a susceptible line during infection with *S. hermonthica* (34) and is associated with increased lignin deposition and post-attachment resistance (35).

**Figure 3:**
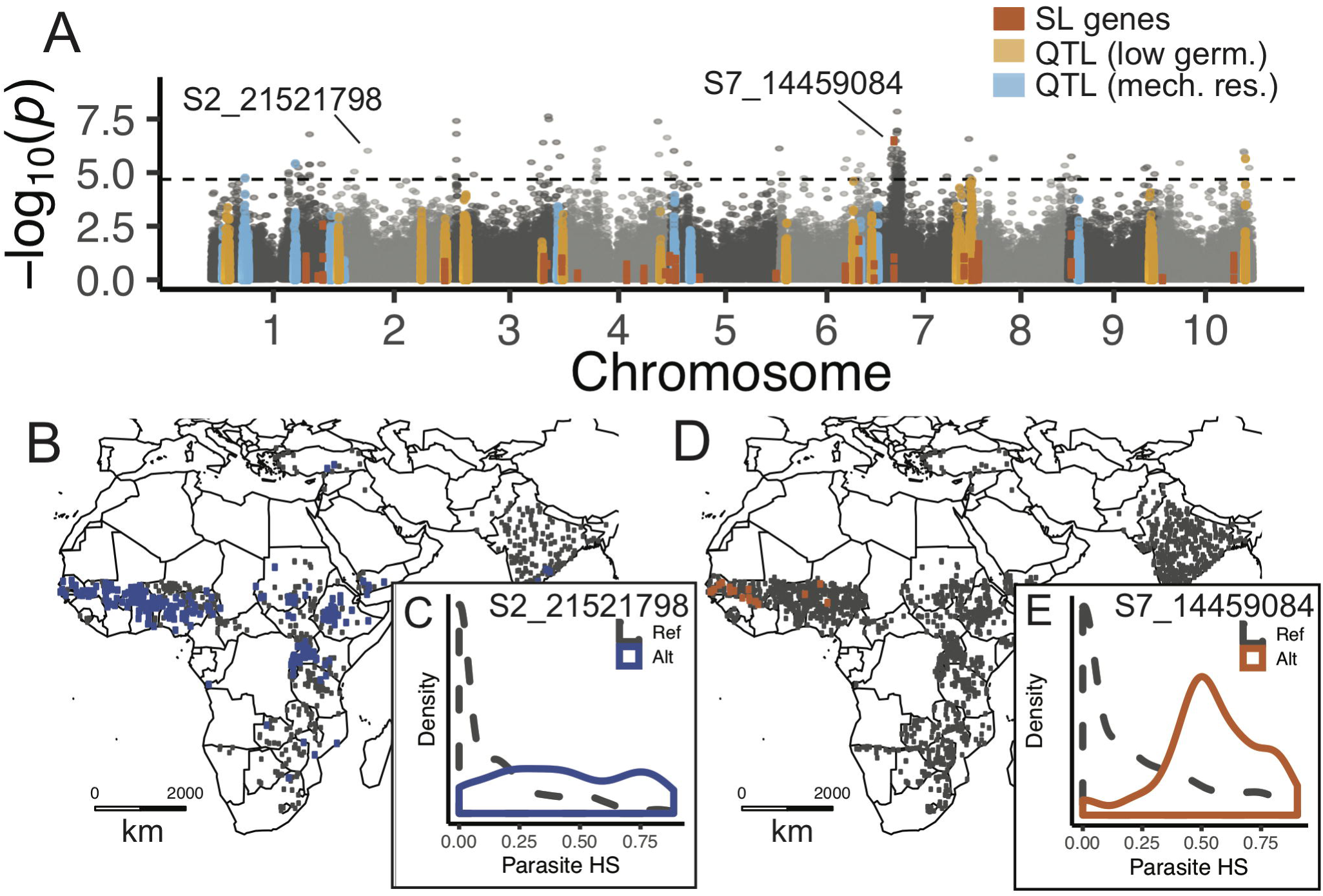
Sorghum genome-wide associations with parasite distribution implicate cell wall and strigolactone signaling genes. A) Genome-wide association with parasite habitat suitability (HS) score, based on 317,294 SNPs with minor allele frequency (MAF) > 0.01 in 2070 sorghum landraces. SNPs in genomic regions linked to strigolactone (SL) biosynthesis/signaling (red), resistance through formation of a mechanical barrier (light blue), or low *Striga hermonthica* germination (orange) are indicated. The dashed line represents significance threshold at a false discovery rate of 5%. B) Geographic distribution of reference and alternate alleles for a SNP (S2_21521798) in a pectinesterase gene (MAF = 0.275). C) Distribution of parasite habitat suitability scores for sorghum accessions segregating for S2_21521798. D) Geographic distribution of reference and alternate alleles for a SNP (S7_14459084) in a gene homologous to SMAX1 (MAF = 0.014). E) Distribution of parasite HS scores for sorghum accessions segregating for S7_14459084.

Across all gene models tagged by SNPs in the genome-wide analysis (i.e. within 1 kb), no GO term met the threshold for significance after correction for multiple comparisons. The strongest enrichment scores were in genes with GO terms related to cell wall organization (GO:0071555; corrected *p* = 0.13, mean *p* score for 48 genes = 0.27), cell wall (GO:0005618, corrected *p* = 0.17, mean *p* score for 70 genes = 0.29), and pectinesterase activity (GO:0030599, corrected *p* = 0.22, mean *p* score for 39 genes = 0.26). The strongest SNP association to parasite occurrence in a pectinesterase gene model was in *Sobic.002G138400* (SNP S2_21521798, *p =* 0.01; Table S4). Although the allele associated with parasite occurrence at S2_21521798 is not predicted to cause an amino acid change, it tagged a haplotype block encompassing complex structural variation 204.4 kb upstream of the gene model, suggesting a potential regulatory variant or nearby presence/absence variation. Overall, SNPs in genes related to strigolactone biosynthesis and signaling (Table S3) did not show a significant enrichment for associations with *S. hermonthica* distribution (uncorrected *p* = 0.09).

### Signatures of balancing selection in candidate regions

We further investigated three candidate genes with polymorphism that exhibited distinct geographic patterns and had known or potential roles in *S. hermonthica* resistance. Elevated Tajima’s D values can indicate an excess of shared polymorphism at a locus, expected for regions of the genome under balancing selection, whereas strongly negative values can indicate an excess of low frequency polymorphism, expected under either purifying or positive selection. Two 5 kb genomic regions, spanning SNPs in *LGS1* (SNP S5_69986146 in gene model *Sobic.G005G213600*) and a pectinesterase gene (SNP S2_21521798 in gene model *Sobic.002G138400*, MAF=0.275) exhibited elevated values of Tajima’s D compared to 1000 randomly sampled 5 kb windows containing or overlapping gene models (Fig. S6; *p* = 0.02, *Sobic.G005G213600*; *p =* 0.05, *Sobic.002G138400*). Regions of elevated Tajima’s D were localized to relatively small windows centered on SNPs associated with *S. hermonthica* habitat suitability, and larger window sizes produced weaker signals. We looked for these alleles in several genomes of wild relatives of sorghum but found no reads mapping to the pectinesterase gene and no evidence for the *LGS1* loss-of-function alleles characterized using data from previously sequenced accessions of *S. propinquum* (Kunth) Hitchc., *n*=2, and *S. arundinaceum* (as synonym *S. bicolor* subsp. *Verticilliflorum* (Steud.) de Wet ex Wiersema & J.Dahlb., *n*=2)(36).

We did not observe strong departures from the neutral expectation of *D* for the region surrounding a gene with homology to *SMAX1* (gene model *Sobic.007G090900* tagged by SNP S7_14459084, *p*=0.6). The minor allele at S7_14459084 was at low frequency in the GBS dataset (MAF = 0.014) and most common in West Africa (Fig. 3C), which is not well sampled in the WGS dataset. The signal of association with *S. hermonthica* occurrence extended more than 7.5 Mb on Chromosome 7 (Fig. 3A), but we did not have sufficient sampling to suggest that S7_14459084 tags an incomplete or soft sweep in either the GBS or WGS datasets according to the haplotype-based statistic *n*S_L_ (37).

### LGS1*-mediated resistance depends on parasite population and environment*

We further characterized the effects of loss-of-function variation at *LGS1* by comparing multiple aspects of performance of newly generated lines of sorghum with CRISPR-Cas9 deletions of *LGS1*. Root exudate from these *LGS1* deletion lines induced substantially lower *S. hermonthica* germination compared to control lines (*p* < 0.001, likelihood ratio test for fixed effect of deletion; Fig. 4A). However, the benefit of *LGS1* deletion was conditional on the specific parasite population (*p*=0.005; likelihood ratio test). We observed <1% germination in response to exudate from *LGS1* deletion lines in an *S. hermonthica* population from Siby, Mali (95% Wald CI: 0.0, 4.0% germination) under the most stressful treatment (drought and low nutrient) but 6.1% germination in an *S. hermonthica* population from Kibos, Kenya (95% Wald CI: 0.2, 12.4% germination). In contrast, exudate from wild-type Macia grown under the same conditions induced mean germination of 40% (Siby; 78% germination in response to 0.1 ppm GR24) or 29% (Kibos; 66% germination in response to 0.1 ppm GR24). The Kenyan *S. hermonthica* similarly showed higher germination sensitivity to an orobanchol standard compared to the Malian *S. hermonthica* (Fig. 4C). The fact that Kenyan *S. hermonthica* showed similar germination response to 5-deoxystrigol and orobanchol standards, while germinating at a higher rate on intact *LGS1* alleles vs. deletions, suggests that *LGS1* deletion has effects on SLs in exudate beyond changing the ratio of 5-deoxystrigol to orobanchol in exudate.

**Figure 4.**
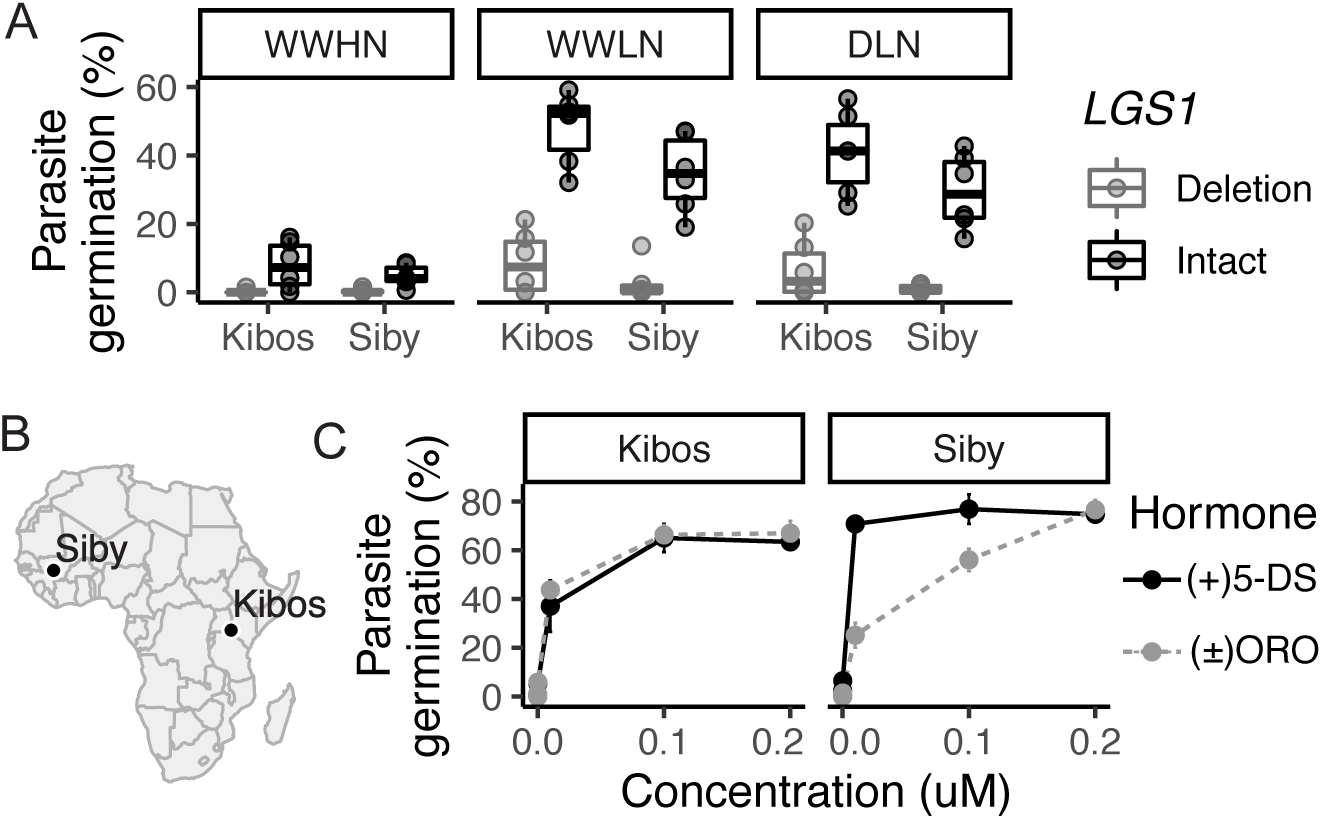
Efficacy of *LGS1* loss-of-function varies across abiotic and biotic gradients. A) Germination of two *S. hermonthica* populations in response to root exudate from 3-wk-old wild-type Macia and CRISPR-Cas9 edited sorghum seedlings with *lgs1* deletion alleles. Plants were grown under well-watered high nutrient (WWHN), well-watered low nutrient (WWLN), or drought low-nutrient (DLN) conditions. Germination was 0% in response to diH_2_O and 66% (Kibos) or 78% (Siby) in response to 0.1ppm GR24. B) Origin of two tested *S. hermonthica* populations. C) Germination of two *S. hermonthica* populations in response to synthetic strigolactones, (+)5-deoxystrigol (5-DS) or (±)orobanchol (ORO).

### LGS1 *loss-of-function reduces expression of photosystem genes*

Changes in strigolactones (and indeed any hormone) are potentially pleiotropic given their many functions, suggesting trade-offs may be associated with *LGS1* variation. In CRISPR-Cas9 edited sorghum, we found 505 differentially expressed genes in roots (244 downregulated in *LGS1* knockout lines and 261 upregulated) or 2167 differentially expressed in shoot (917 downregulated in *LGS1* knockout lines and 1250 upregulated) at 6 weeks after planting compared to wild-type Macia. Of all differentially expressed genes, 185 were differentially regulated in both root and shoot, including a transcription factor most highly expressed in sorghum panicles during floral initiation (*Sobic.010G180200*, downregulated in knockout) and a GRAS family transcription factor homologous to *RGA*, which in *A. thaliana* represses giberellic acid-induced vegetative growth and floral initiation (*Sobic.008G168400*, upregulated in knockout) (38). In shoots, photosystem II genes were most enriched among differentially expressed genes (GO:0009523, *p*<0.001; 11/36 genes with corrected *p* < 0.05; Supplemental Data File S2). Only GO-terms related to photosystem I (GO:0009522, *p <* 0.001; 8/10 genes with corrected *p <*0.1) and photosynthetic light harvesting (GO:0009765, *p <* 0.001; 12/18 genes with corrected *p* <0.05) were enriched among differentially expressed genes in roots. All differentially expressed photosystem I, photosystem II, and light-harvesting genes were downregulated in *LGS1* knockout lines, with lower expression in roots than shoots, and 1.7 to 7.2x higher expression in Macia wild-type (log_2_ fold-change ratios ranging from 0.77 to 2.84; Supplemental Data File S2).

Six of the light-harvesting or photosystem I genes downregulated in roots of *LGS1* knockout lines were also differentially expressed in nutrient-stressed roots of a resistant sorghum line (SRN39), which produces high levels of orobanchol, compared to a susceptible sorghum line (Shanqui Red) that produces mainly 5-deoxystrigol (29) (Supplemental Data File S2). Strigolactone biosynthesis genes were also upregulated in the *LGS1-*deficient sorghum line SRN39 compared to the susceptible line, including two genes with homology to *NSP1* (*Sobic.001G341400*, *p* < 0.001 and *Sobic.002G372100*, *p =* 0.03), *CCD7* (*Sobic.006G170300*, *p* <0.001), and *LBO* (*Sobic.003G418000*, *p* = 0.01; Fig. S7, Table S3). Of these four genes, only *LBO* was also significantly upregulated in the CRISPR *LGS1* knockout line (in shoots), suggesting the natural low germination stimulating line carries additional alleles altering SL profiles.

In addition to reduced expression of photosynthesis genes, we found evidence for subtle effects of *LGS1* loss-of-function on sorghum growth. Total leaf area of 65-day-old plants was not significantly different among three independent CRISPR knockout lines grown in nutrient rich soil (Fig. S8). In contrast, deletion lines had smaller leaf area relative to wild-type control and two event-null lines (*p <* 0.05, Scheffé’s method, Fig. S8). However, no significant differences were seen in a separate experiment on younger plants grown under nutrient limitation, in which we did not detect a significant effect of *LGS1* deletion on dry biomass, specific leaf area, SPAD-based measurement of leaf chlorophyll, or root number. Consistent with differential expression of transcription factors involved in growth and floral initiation (e.g. *Sobic.008G168400*), it may be that phenotypic differences are only detectable later in development or in some environments.

## DISCUSSION

Pests, pathogens, and parasites threaten human health and food security in a changing world but understanding mechanisms of resistance across diverse taxa remains challenging. Here, we evaluate the hypothesis that geographic selection mosaics maintain genetic diversity in host resistance alleles across gradients of parasite occurrence in smallholder farming systems. We extended genotype-environment association analyses to biotic environmental gradients using species distribution models to model high resolution variation in parasite occurrence at continent scales. We report strong associations with parasite occurrence for novel candidate resistance loci in the sorghum-*Striga hermonthica* pathosystem and characterize diverse loss-of-function mutations in the sorghum resistance gene *LGS1*. Geographic distribution of loss-of-function alleles suggests that *LGS1-*conferred resistance is stable across some range of environments and genetic backgrounds. However, the low frequency and paucity of *LGS1* loss-of-function alleles outside of parasite-prone areas, combined with the germination sensitivity of some parasite populations to orobanchol and impacts on host photosynthesis regulation, suggest there may be trade-offs associated with *LGS1* loss-of-function.

Our results support the hypothesis that spatial variation in selective pressures controls geographic clines in the frequency of host resistance alleles. The patterns we characterized are likely representative of long-term averaged conditions as opposed to a snapshot of coevolution because our parasite distribution model used occurrence records spanning more than 150 years, and the landraces we studied were collected across the last several decades. In addition to temporal genetic variation (for example, due to negative frequency-dependent selection), many parasites like *S. hermonthica* are highly genetically diverse across space (26), so that host resistance phenotype depends on local parasite genotypes (39). Depending on the parasite genotype used in experiments, this host by parasite genotype interaction might obscure GWAS for host resistance (40). By capturing variation at coarse scales, the spatial perspective of parasite-associated host genomic variation presented here could facilitate complementary, inexpensive detection of genomic regions contributing to resistance across diverse parasites.

Our study revealed evidence for locally-adaptive natural variation in genes related to cell wall modification. Cell wall modifying enzymes including pectinesterases are highly expressed in the developing haustorium of parasitic plants and in the host-parasite interface (41–44). Pectinesterases de-esterify pectin in plant cell walls, making it accessible to other cell wall degrading enzymes and loosening cell walls. However, some studies have suggested that in the presence of Ca^2+^, de-esterified pectin forms egg-box structures and instead stiffens cell walls (45). Rigidification of sorghum cell walls by their own pectinesterases (such as *Sobic.002G138400*, Table S4) or reduced activity could help defend against parasitic invaders. Notably, Yang *et al.* (2015) reported haustorium specific expression and positive selection on pectinesterase inhibitors in parasitic plant lineages and a pectinesterase inhibitor showed exceptionally high host species-associated expression in field populations of *S. hermonthica* (46). Parasite inhibitors could interact with host pectinesterases or help maintain integrity of parasite cell walls in the face of high pectinesterase expression during haustorial invasion.

Our results also suggest that *LGS1* loss-of-function alleles may be adaptive in *S. hermonthica-*prone regions, but that costs of resistance may limit their distribution.

Loss-of-function alleles are relatively uncommon but higher in frequency and broadly distributed where parasites occur (Fig. 2C). Associations for the putative resistance locus *LGS1* were not statistically significant after correcting for kinship, likely due to covariation with genomic background (Fig. S5), which has been shown to substantially reduce power to detect causal loci in locally-adapted sorghum landraces (14) and in simulations (47). Experiments with genome-edited sorghum, however, supported an adaptive role of *LGS1* loss-of-function, particularly in marginal environments (Fig. 4A). The diversity of loss-of-function variants reported here (Fig. 2A-B) and elsewhere (29), their wide geographic distribution (Fig. 2C), and an excess of high frequency polymorphism localized to *LGS1* (Fig. 4) are further consistent with long-term maintenance of *LGS1* diversity under balancing selection. The underlying mechanisms could include negative frequency dependent selection or spatiotemporally variable selection favoring different alleles in different environments, depending on the relative costs of resistance.

Costs of resistance linked to impacts of SL structural changes on host fitness could include impaired signaling to AM fungi, susceptibility to *S. hermonthica* genotypes sensitive to orobanchol, or impacts on endogenous strigolactone signaling. Consistent with the second hypothesis, we found higher sensitivity to root exudates from CRISPR-edited *LGS1* deletion lines for *S. hermonthica* from Kenya compared to *S. hermonthica* from Mali (Fig. 4). Our findings indicate the existence of *S. hermonthica* alleles that promote germination on orobanchol, alleles that could increase in frequency following an increase in cultivation of *LGS1* deletion sorghum. Consistent with the third hypothesis, we found evidence of systemic downregulation of photosystem-related genes in *LGS1* knockout lines, corresponding to a previously identified role of SLs as positive regulators of light harvesting (48). Perturbations of photosynthesis regulatory networks due to changes in strigolactone profiles could be buffered by other components of the system, ultimately resulting in subtle but detectable impacts on host fitness (Fig. S8).

Taken together, this study provides evidence of locally-adaptive natural variation in sorghum parasite resistance genes across African smallholder farming systems. We describe long-term maintenance of diversity in known and novel candidates implicated in pre- and post-attachment resistance to the parasitic plant *Striga hermonthica*. However, the possibility of trade-offs and the existence of orobanchol-sensitive *S. hermonthica* populations suggest potential pitfalls with widespread deployment of the *LGS1* loss-of function allele in sorghum cultivation, particularly under low nutrient conditions and in fields with high parasite density. Our findings highlight the complexity of interacting abiotic, biotic, and human pressures shaping genome polymorphism across environments in cultivated species.

## METHODS

### Species distribution models

Genome-environment association analyses are designed to identify putatively locally adaptive genetic loci where allelic variation is strongly associated with home environments (49). To employ this approach with biotic gradients, we required information on local parasite pressure for each sorghum landrace. We used species distribution models (SDMs) to estimate habitat suitability of *Striga hermonthica* at the location of each georeferenced sorghum landrace, under the assumption that modeled habitat suitability scores are a reasonable proxy of parasite success averaged over the long term and in comparison with sites where the parasite never occurs.

*Striga hermonthica* SDMs were constructed with Maxent, a machine learning tool for predicting habitat suitability for a species of interest given a set of environmental variables and presence-only data (31). We compiled 1369 occurrence records for *S. hermonthica* (Supplemental Data File S3; additional details in Supplemental Materials & Methods). We also created ‘sorghum-only’ models based on a subset of *S. hermonthica* records (*n* = 262) that were annotated as occurring specifically on sorghum. Environmental variables were chosen based on prior knowledge of the ecology of *S. hermonthica* (50). We included bioclimatic and topographic variables (annual rainfall, mean temperature of the wettest quarter, isothermality, potential evapotranspiration [PET], and topographic wetness index) from CHELSA (51) and ENVIREM datasets (52). Soil variables (clay content, nitrogen, and phosphorus) were based on continental and global-scale soil property maps (53, 54). SDMs were implemented and evaluated with ENMeval, using the ‘checkerboard2’ method for data partitioning, which is designed to reduce spatial autocorrelation between testing and training records (55).

### *Sorghum* LGS1 *loss-of-function alleles*

Fine-scale natural variation in sorghum *LGS1* was characterized using whole genome sequencing (WGS) data from a set of 143 georeferenced landraces from the sorghum bioenergy association panel (BAP) (56). The BAP includes both sweet and biomass sorghum types, accessions from the five major sorghum botanical races (durra, caudatum, bicolor, guinea, and kafir), and accessions from Africa, Asia, and the Americas. The BAP accessions were sequenced to approximately 25x coverage and genotyped as part of the TERRA-REF project (www.terraref.org). This whole genome sequencing derived dataset is referred to throughout the manuscript as the ‘WGS dataset’ to distinguish it from the genotyping-by-sequencing dataset used for GEA.

We characterized three loss-of-function alleles in *LGS1* using data from the WGS dataset. Frameshift and nonsense mutations were identified using *SnpEff* v4.3t for SNP calls and small indels in *Sobic.005G213600* (57). To characterize large deletion variants, we quality trimmed reads with BBDuk (sourceforge.net/projects/bbmap/; qtrim=rl, trimq=20) and aligned to the *Sorghum bicolor* v3.1 reference genome (DOE-JGI, http://phytozome.jgi.doe.gov/) with BWA MEM v0.7.17 (58). Duplicates were removed with SAMBLASTER v0.1.24 (59) and structural variants were called for each landrace with LUMPY v0.2.13 (60). SVTYPER v0.6.0 was used to call genotypes for structural variants ≤ 1 Mb spanning *Sobic.005G213600* (61).

After characterizing the *LGS1* deletion breakpoints using the WGS dataset, we imputed deletion calls to the GBS dataset. We considered the *LGS1* region to be deleted if at least one SNP was called in the 5 kb region flanking positions of deletion breakpoints, but all data were missing between breakpoints. We considered the *LGS1* region to be present if at least one SNP was called within the *Sobic.005G213600* gene model. Fifteen low-coverage samples, with missing data extending 5 kb into flanking regions, were excluded.

### *Experimental validation of* LGS1 *deletion and frameshift alleles*

*LGS1* loss-of-function alleles were validated by testing 12 accessions from the Sorghum Association Panel (SAP) (62) for their ability to stimulate *S. hermonthica* germination (Table S5). Seed was obtained from the U.S. National Plant Germplasm System (NGPS) through GRIN (USDA, ARS, PGRCU). Two accessions were previously reported to be resistant to *S. hermonthica* due to low germination stimulation, two accessions were susceptible, and eight accessions had unknown resistance (29, 63). Root exudates were harvested 43 days after planting and used for *S. hermonthica* germination assays (see Supplemental Materials & Methods for a detailed description of plant growth conditions and germination assays).

### Genome-environment associations

We performed a genome-wide scan for SNPs in the sorghum genome strongly associated with values of habitat suitability estimated by our *S. hermonthica* distribution model. Sorghum genotypic information was extracted from a public dataset of accessions genotyped using genotyping-by-sequencing (GBS) (14, 64–66). This dataset comprises a diverse set of worldwide accessions including germplasm from the SAP (62), the Mini-Core Collection (67), and the Bioenergy Association Panel (BAP) (56). Beagle 4.1 was used to impute missing data based on the Li and Stephens (2003) haplotype frequency model (68). The average missing rate in the non-imputed dataset is 0.39 (66). After excluding sorghum accessions with missing coordinates and SNPs with minor allele frequency less than 0.01, the dataset, referred to throughout the manuscript as the ‘GBS dataset’, included 1547 African landraces among 2070 georeferenced accessions total genotyped at 317,294 SNPs.

At each location of a georeferenced accession in the GBS dataset, we extracted logistic output from the *S. hermonthica* distribution model (Supplemental Data File S4-5) as the environment. To account for regions where predicted habitat suitability is high but *S. hermonthica* has not been recorded, we cropped model predictions to within 200 km from any occurrence record and set values outside of this range to zero to derive for each grid cell an *S. hermonthica* occurrence score ranging from zero to one; more than half of sorghum accessions are from locations with parasite HS scores greater than zero (Fig. 1B). Genome-wide associations for each SNP with *S. hermonthica* occurrence were computed for the GBS dataset using a mixed linear model (MLM) fit with GEMMA v0.94 (69). To take into account relatedness among individuals, we used a centered kinship matrix (-gk 1) generated from all 317,294 SNPs before calculation of p score statistics (-lmm 3). P scores were adjusted for multiple comparisons using the Benjamini and Hochberg (1995) procedure (FDR = 0.05). To visualize genomic regions previously implicated in resistance to *S. hermonthica*, locations of QTL from the linkage mapping study of Haussman et al. (2004) in the *S. bicolor* v3.0 genome were downloaded from the Sorghum QTL Atlas (http://aussorgm.org.au) (70). We tested for associations with *LGS1* loss-of-function mutations using the same procedure and kinship matrix as for the genome-wide association analysis.

We identified gene functions enriched for associations with parasite distribution using the gene score resampling method in ErmineJ (71). This method places higher value on gene scores than their relative ranking and does not require choice of a threshold for significance. For each gene model, we used the lowest p score from GEMMA of any SNP within 1 kb of gene model boundaries, and enrichment analyses were performed using the mean of all gene scores in the gene set. Gene sets were created using GO terms for all gene models in the *Sorghum bicolor* v3.0 genome (annotation version 3.1). We also created a custom gene set comprising 30 gene models implicated in strigolactone biosynthesis and signaling (Table S3). Enrichment analysis was performed with 200,000 iterations, excluding gene sets with less than 5 or more than 200 genes.

### Signatures of selection in candidate genomic regions

We performed scans for selection in 1 Mb regions surrounding focal SNPs. Linkage disequilibrium between sites was determined with vcftools v0.1.15 (--geno-r2 parameter). To identify regions under balancing selection among a subset of African landraces, we used Tajima’s D calculated with vcftools in non-overlapping 5 kb windows, excluding SNPs with more than 70% missing data. *P-*values for candidate regions under selection were calculated based on the empirical distribution of Tajima’s D for 1000 randomly sampled 5 kb windows that overlapped or fully encompassed gene models. We searched for sweeps using the *n*S_L_ statistic with selscan v1.2.0a (72).

### Performance of CRISPR-Cas9 edited sorghum

We evaluated potential trade-offs to *LGS1* loss-of-function by testing *S. hermonthica* germination response and growth of genome edited sorghum. *LGS1* was deleted in the sorghum line Macia using the CRISPR-Cas9 system to produce three independent genome-edited lines, with sequence confirmation using Southern-by-Sequencing (see Supplemental Materials & Methods).

We measured plant growth rate in a greenhouse at Corteva Agriscience using a hyperspectal imaging system to determine total leaf area of each plant (day temperature: 26°C; night temperature: 20°C; 16 hr photoperiod). We compared three independent CRISPR-edited lines (*lgs1-d* lines 1-3), the wild-type Macia line, and two event-null transformation lines that had intact *LGS1* but SNPs at the two cutting sites flanking *LGS1*. Experimental design was unbalanced incomplete block with replications of 10 to 33 plants. Images were collected weekly on days 23-65 after planting. Total leaf area per plant was analyzed using one-way ANOVA followed by Scheffé’s method.

In a separate experiment at Penn State, we compared the same three CRISPR-edited lines and two independently bulked seed batches of wild-type Macia under three conditions: well-watered with high nutrition (WWHN), well-watered with low nutrition (WWLN), and drought with low nutrition (DLN). Plants (*n =* 19 per line and treatment) were grown in autoclaved sand in cone-tainers under natural light conditions in a greenhouse (day temperature: 27°C; night temperature: 24°C). For the WWHN treatment, plants were fertilized every other day with 15 mL of 1/10^th^ strength Miracle-Gro Plant Food (24% nitrogen, 8% phosphorus, 16% potassium) and watered with 15 ml of tap water on days without fertilization. Plants were watered every day with 15 mL tap water (WWLN treatment) or every four days with 15 mL water (DLN treatment). Three plants per line and treatment were harvested at 21 DAP for collection of root exudates as described for validation of natural deletion alleles (see Supplemental Materials & Methods). Germination assays testing root exudate in 12-well plates (three technical replicates per biological replicate, ~60 seeds per well) were performed in the USDA quarantine facility at Penn State, using *S. hermonthica* seeds collected in 2016 from a sorghum field in Kibos, Kenya (0°40’S; 34°49’E) and in 2018 from a sorghum field in Siby, Mali (12°23’ N; 8°20’ W). Germination assays used a preconditioning temperature of 29°C for 11 days in 1 mL of diH_2_O prior to addition of 1.5 mL root exudate. A complementary assay tested the response of the same *S. hermonthica* populations to five concentrations (0, 1×10^−10^, 1×10^−8^, 1×10^−7^, and 2×10^−7^ M) of (±)orobanchol and (+)5-deoxystrigol (CAS 220493-64-1 and 151716-18-6; Olchemim, Czech Republic).

To compare performance of wild-type and CRISPR-edited sorghum, we also measured biomass (shoot and root dried weight), chlorophyll concentration (measured via SPAD meter), root architecture traits (number of crown, adventitious [shoot-borne], and seminal roots), shoot architecture traits (plant height, leaf number, total shoot length, and leaf internode lengths), and specific leaf area (SLA) at 41 DAP. We tested significance of deletion alleles using generalized linear mixed models for count data (root number) or linear mixed effects models (all other traits) in the R package lme4 (73), where deletion and treatment were fixed effects and sorghum lines were random effects. For germination rate data, fixed effects of *S. hermonthica* population (Kibos or Siby) and root weight were also included.

### LGS1 and expression of genes in strigolactone synthesis and signaling pathways

To assess the impact of putatively locally adaptive variation at *LGS1*, we studied transcriptomes for roots and shoots (excluding leaf blade) of CRISPR knock-out (*lgs1-d* line 3) and wild-type Macia lines at 30 days after planting under WWLN, the treatment under which plants exuded the most SLs (Fig. 4). RNA extractions were performed with the NucleoSpin RNA Plant kit (Machery-Nagel). We used the 3’-TagSeq approach (74) which quantifies mRNA based on the 3’ end of transcripts. cDNA libraries were sequenced on a shared lane of an Illumina Nova-Seq at the Genomic Sequencing and Analysis Facility at the University of Texas at Austin. We recovered between 10 and 15 million raw single-end 100 bp reads per sample and compared expression differences using the TagSeq v2.0 pipeline (https://github.com/Eli-Meyer/TagSeq_utilities/). We also generated TagSeq libraries using root tissue from five replicate individuals each of sorghum lines Shanqui Red (PI 656025) and SRN39 (PI 656027) grown under nutrient deficient conditions (see Supplemental Materials & Methods). Count data were analyzed in DESeq2 with FDR correction (□=0.05) (75). Enrichment analysis was performed as for genome-wide association analysis, except only the set of gene models with non-zero expression were used as the background.

## Data Availability

Raw reads generated for the Tag-Seq study have been deposited in the NCBI SRA database under BioProject accession PRJNA542394. *S. hermonthica* occurrence records and species distribution models are available in the supporting information.

## Acknowledgments

We thank the many collectors, volunteers, and herbarium curators who made this work possible and are particularly grateful to Marie-Hélène Weech and the staff of the Royal Botanic Gardens Kew and the Muséum national d’Histoire naturelle, Paris. Historical data from French herbaria were obtained thanks to “Les Herbonautes” (MNHN/Tela Botanica), part of Infrastructure Nationale e-RECOLNAT (ANR-11-INBS-0004). We thank Alice MacQueen for comments that improved the manuscript, Ping Che for assistance with tissue culture, Meizhu Yang for molecular analysis of CRISPR mutants, Eric Schultz for hyperspectral phenotyping, and the Dean of Eberly College and Head of the Department of Biology for their support in construction of the USDA-APHIS certified Parasitic Plant Containment Laboratory at Penn State. Whole genome sequence data used here is from the TERRA-REF project, funded by the Advanced Research Projects Agency-Energy (ARPA-E), U.S. Department of Energy, under Award Number DE-AR0000594. This study is based on work supported by a National Science Foundation Postdoctoral Research Fellowship in Biology to ESB under Grant No. 1711950. The views and opinions of authors expressed herein do not necessarily state or reflect those of the United States Government or any agency thereof.

## Notes

#### Summary of Updates

We have included additional experiments testing impact of LGS1 loss-of-function using genome-edited sorghum lines.

