## Supplementary Appendix for "Genomics of sorghum local adaptation to a parasitic plant"

### Supplementary Information Text

**Detailed Materials and Methods.** This section contains detailed description of methods for plant growth conditions, species distribution modeling, generation of CRISPR-Cas9 edited sorghum mutants, preparation and analysis of TagSeq libraries, and references for *S. hermonthica* occurrence records from published studies.

#### Collection of sorghum root exudates to validate imputation of *LGS1* deletions from GBS dataset.

Accessions were grown individually in two-gallon pots with 70% potting mix (Premier pro-mix PGX) and 30% medium commercial grade sand in a greenhouse at 85°F during the day and 75°F at night with a 16-hour photoperiod. Pots were fertilized with 1x strength Miracle-Gro (Miracle-Gro® Water-Soluble All-Purpose Plant Food, The Scotts Company, LLC., Marysville, OH) once at fourteen days after planting. Forty-three days after planting, potting components were carefully washed from roots and whole plants were placed in separate flasks with a 1:5 ratio of root:DI water (v/v). Flasks were sealed with parafilm to prevent evaporation and placed into darkness at room temperature. After 48 hours, root exudate from each plant was centrifuged at 9,000 rpm for 10 minutes before removing supernatant for germination assays.

Germination trials were conducted using seed of *Striga hermonthica* collected on sorghum in 2013 from Alupe, Kenya and assayed in the USDA quarantine lab at the University of Virginia. Seeds were surface sterilized with 0.5% bleach, rinsed three times with sterile distilled water, and 75 seeds were transferred with 500uL di$H_2O$ to 12-well microtiter plates. After 10 days of preconditioning in the dark at 30ºC, 2.25 mL of fresh root exudate was applied to each well. Exudate from five biological replicates (sorghum individuals) per sorghum genotype, with three technical replicates (wells) per biological replicate, were tested. GR24 at two concentrations, 1 ppm and 0.1 ppm, and water were used as positive and negative controls. Germinated seeds were counted under a stereomicroscope after 66 hours incubation. We tested significance of deletion alleles using linear mixed effects models in the R package lme4, where deletion was a fixed effect and genotypes were random effects.

#### Species distribution models.

*Striga hermonthica* SDMs were constructed with Maxent, a machine learning tool for predicting habitat suitability for a species of interest given a set of environmental variables and presence-only data (1). We compiled 1369 occurrence records for *S. hermonthica* records downloaded from the Global Biodiversity Information Facility (www.gbif.org, https://doi.org/10.15468/dl.mvzauk), newly digitized records for herbarium specimens housed in the collections of the Royal Botanic Gardens Kew (collection: K), the National Museum of Natural History in Paris (MNHN; collection: P), the French Agricultural Research Centre for International Development (CIRAD; collection ALF), the University of Montpellier (collection MPU), and the Botanical Garden of Lyon (collection LYJB), and observations from published studies (2–17) (Supplemental Data File S3). Records within 0.01 degree (~1 km) of another observation were excluded to reduce sampling bias. To characterize the background environment across the study extent, we sampled 10,000 points at random from a 500 km radius surrounding locations of 1,050 occurrences in the final dataset. We also created 'sorghum-only' models based on a subset of *S. hermonthica* records ($n$ = 262) that were annotated as occurring specifically on sorghum.

Environmental variables were chosen based on prior knowledge of the ecology of *S. hermonthica* (2). Bioclimatic and topographic variables (annual rainfall, mean temperature of the wettest quarter, isothermality, potential evapotranspiration [PET], and topographic wetness index) were obtained from CHELSA (18) and ENVIREM datasets (19). Soil variables (clay content, nitrogen, and phosphorus) were based on continental and global-scale soil property maps (20, 21). We explored additional ecologically relevant variables but did not include them in the final model due to high correlation across the study background with annual rainfall (correlated with soil PH, aluminum, and precipitation seasonality) or soil clay content (correlated with sand fraction) as indicated by Pearson coefficients ($|r| > 0.7$).

SDMs were implemented and evaluated with ENMeval, using the 'checkerboard2' method for data partitioning, which is designed to reduce spatial autocorrelation between testing and training records (22). The distribution model with the lowest AICc was selected for further comparisons with genome variation in sorghum. Two niche overlap statistics, Schoener's $D$ (23) and the $I$ similarity statistic (24) were calculated for the all-occurrence and sorghum-only models using the R package *dismo* (25).

**CRISPR-Cas9 editing of sorghum *LGS1*: Plasmid construction.**
To delete the *LGS1* gene in the sorghum line Macia using the CRISPR-Cas9 system, two single-guide RNAs (sgRNAs), designated as CR1 and CR2 were designed to recognize two CRISPR target sites TS1 and TS2 (Figure 7a). The CR1/CR2 pair was used to delete the complete *LGS1* coding sequence and a portion of the promoter.

Plasmids were designed to express single guide RNAs (sgRNA) and the *Streptococcus pyogenes* Cas9 endonuclease. The sgRNA gene was expressed by a maize U6 polymerase III promoter (26). The Cas9 expression cassette contains the maize ubiquitin1 promoter (ubiZM1) and potato protease inhibitor II (*pin*II) terminator. The Cas9 sequence was maize codon optimized, and the potato ST-LS1 intron and the nuclear localization signals from the Simian Virus 40 (SV40) were added for appropriate expression and nuclear targeting in maize, as previously described (26). Constructs were assembled using chemically synthesized DNA fragments with standard DNA techniques. Neomycin phosphotransferase II (NPTII) served as a transformation selection marker. To facilitate delivery of the genome editing reagents (Cas9, guide RNAs) into sorghum cells and regeneration of plants, morphogenic transcription factors *Baby boom* (*Bbm*) and *Wuschel2* (*Wus2*) were expressed under control of the maize *phospholipid transferase* promoter (Zm-PLTPpro) and *auxin-responsive gene1* (Zm-Axig1pro), respectively, and the plasmids were constructed as described (27). The plasmid sequence was deposited in GenBank with accession number of MN585216.

**CRISPR-Cas9 editing of sorghum *LGS1*: Sorghum transformation.**

The genome editing reagents, including CR1, CR2 and *S. pyogenes* Cas9 were introduced into immature embryos via agrobacterium transformation. Agrobacterium containing the sgRNA CR1 and CR2, spCas9, NPTII, BbM and Wus2 gene cassettes (Supplementary Figure S9) were introduced to sorghum Macia immature embryos as described by Che *et al.* (28). Regenerated T0 plants were screened using a PCR assay designed to amplify the rejoined junction of the *LGS1* deletion (Figure S9a, b). Briefly, genomic DNA was extracted from leaf tissues of T0 plants as described in Gao *et al.* (29). PCR was performed using REDTaq ReadyMix (Millipore Sigma) with the primers p1 (5'- GGTATGTCCCTGAGCATGTCT) and p2 (5' - TTGTTTAATTCTTTCATGTGGTTCTATTTGT). Amplicons of ~656 bp are expected when the TS1/TS2 cleavage and NHEJ result in deletion of a 2.9-kb fragment. Next-Generation Sequencing (NGS) was used to evaluate the junction sequence of deletion variants. The junction was amplified first with PHUSION® Flash High Fidelity PCR Master Mix (F-548; Thermo Fisher Scientific, Waltham, MA), and the secondary PCR used Phusion Master Mix F-531 (Thermo Fisher Scientific). The Molecular Inversion Probe (MIP) primers used are listed in Supplementary Table S7. The resulting PCR products were purified with a Qiagen PCR purification spin column, DNA concentrations measured with a Hoechst dye-based fluorometric assay, then sequenced directly with NGS.

From 250 infected embryos, 65 T0 plants were regenerated and 39 T0 plants were positive for *LGS1* gene deletion. Sequence analysis confirmed removal of the *LGS1* gene and rejoining of the chromosomal ends created by double-strand DNA breaks (Figure S9c).

**CRISPR-Cas9 editing of sorghum *LGS1*: Production of S2 progeny.**
Three independent T0 plants with identical junction sequence were selected for further analysis. The selected T0 plants were self-pollinated to produce S1 progeny. Thirty-two S1 plants from each line were genotyped using the junction PCR assay to identify individuals that contained *LGS1* deletion (Supplementary Table S6). In each of the three independent lines, the S1 plants segregated at ratio of 3:1 for deletion (*LGS1 lgs1* and *lgs1 lgs1*) and no deletion (*LGS1 LGS1*). S1 plants were also screened for T-DNA integration in the genome using quantitative PCR (qPCR) with primers and probes that amplify the transgenes on T-DNA (Supplementary Table S7, Figure S10). qPCR was performed using Qiagen QuantiTect Multiplex PCR Master Mix. Primers and probes for *Cas9*, *sgRNAs*, *NPTII, Bbm, and Wus2* are listed in the Supplementary Table S8. The S1 plants free (null) of T-DNA were selected for further confirmation with Southern-by-Sequencing (SbS), which can thoroughly detect all the sequences on T-DNA (30). To perform SbS, DNA was extracted from leaf punches of selected S1 *LGS1* deletion plants as described previously. A capture-probe library was created to cover the entire length of the T-DNA used in transformation (Figure S10). Illumina whole genome sequencing libraries were constructed from DNA derived from plants. Hybridizations and sequencing were carried out as described previously (30). Homozygous *lgs1* plants free of T-DNA were selected to produce S2 progeny for phenotyping.

**Root RNA extraction and DNase treatment for SRN39 and Shanqui Red.**

We generated TagSeq libraries using root tissue from five replicate individuals each of sorghum lines Shanqui Red (PI 656025) and SRN39 (PI 656027), grown under nutrient deficient conditions. Shanqui Red is susceptible to *S. hermonthica*, whereas SRN39 is resistant as a result of a ~34 kb deletion spanning *LGS1* and four adjacent genes (29). Accessions were grown individually in two-gallon pots with 70% potting mix (Premier pro-mix PGX) and 30% medium commercial grade sand in a greenhouse at 85°F during the day and 75°F at night with a 16-hour photoperiod. Pots were fertilized with 1x strength Miracle-Gro (Miracle-Gro® Water-Soluble All-Purpose Plant Food, The Scotts Company, LLC., Marysville, OH) once at fourteen days after planting. Twenty-five days after planting, all potting components were carefully washed from roots, and 2-5 ml of the fine root tissue was sampled, placed into labeled vials and stored on dry ice for 1 day during shipment to U of Texas, where samples were stored at -80°C until RNA extraction.

For RNA extraction from sorghum lines Shanqui Red and SRN39, root tissues were ground in liquid nitrogen using mortar and pestle. Ground tissue powder was homogenized using an equal volume (W/V) of TRIzol™ reagent (Invitrogen, Cat#15596018) and 1 mL homogenate was used in RNA extraction. RNA extraction generally followed TRIzol™ Reagent user guide. In brief, 200 µL of Chloroform:Isoamyl alcohol 24:1 (Sigma, C0549) was added to 1 mL homogenate and incubated for 10 min on rotator mixer at room temperature. The homogenate was centrifuged at 12000 RCF for 15 min at 4°C. The clear upper aqueous phase was transferred to new 1.5 mL Eppendorf tubes and combined with an equal volume (V/V) of isopropanol to precipitate nucleic acid. Samples were mixed well by inverting the tubes several times and centrifuged as described above to obtain RNA pellet. The supernatant was discarded, and the pellet washed with 75% alcohol. The pellet was air dried for 10 min at room temperature and suspended in 30 µL of 10 mM Tris-Cl (pH 8.0). RNA samples were left overnight in 4°C refrigerator to dissolve the pellet. Further, RNA samples were treated with DNase I (Ambion™, AM2222) to remove contaminating DNA. In brief, 25 µL of RNA was incubated with 2 units of DNase I and 3 µL of 10X DNaseI buffer for 30 minutes at 37°C using a water bath. The nucleic acid was re-precipitated, air dried and dissolved in 25 µL of 10 mM Tris-Cl (pH 8.0). The RNA was quantified using NanoDrop® ND-1000 Spectrophotometer and ~100 ng of RNA was analyzed by 1% gel electrophoresis to verify that the RNA was intact and free of genomic DNA contamination.

**Library preparation and sequencing for SRN39 and Shanqui Red.**

We used 3'-TagSeq approach (31) with several modifications to construct cDNA libraries. This method focuses on 3' end of transcripts enriched in a size range of 400-500 bp fragments. Briefly, 1 µg of total RNA from each samples were prepared in a volume of 10 µL and incubated with 8 µL of degradation buffer (0.5mM dNTP mix, 1 mM DTT, 1X first strand buffer, 1 µM 3ILL-30TV oligo-dT primer to aim 3' ends of cDNA) at 70°C for 16 min to achieve desire range of RNA fragments (300-600 bp). The entire fragmented RNA was further used in first strand cDNA (FS cDNA) synthesis with RNA oligo primer (S-ILL-swMW) and SMARTScribe Reverse Transcriptase (Clonetech). Then, cDNA was amplified with 16 cycles using Titanium Taq Polymerase, 3ILL and 5ILL primers, and 10 µL of the FS cDNA as template. The amplified cDNA products were purified using NucleoFast PCR Clean-up kit (Machery-Nagel) and quantified using Qubit dsDNA High sensitivity Assay Kit (Invitrogen) on Qubit® 2.0 Fluorometer.

Further, 50 ng of purified cDNA from each sample was individually barcoded with four cycles of barcoding PCR. We used Illumina specific barcodes (ILL-BC) and multiplexed those using Illumina TruSeq universal adapters (TrusSeq_Un) for use on the Illumina platform. An equal volume of barcoded cDNA libraries was pooled together and purified using PureLink Quick Gel

Extraction and PCR Purification Combo kit (Invitrogen). The purified cDNA library pool was separated on 1.5% agarose in 1X TBE buffer (pH 8.2) to excise the fragment in the 400-500 bp size range. The excised gel fragment was purified using a gel extraction kit as mentioned above. The resulting library pool was loaded into a single lane of an Illumina Hiseq-2500 analyzer at the Genomic Sequencing and Analysis Facility at the University of Texas at Austin. We recovered between 3 to 5 million raw single-end 100 bp reads per sample. All primers used in the library preparation are given below.

**Bioinformatic analysis of TagSeq data.**
Briefly, we excluded reads containing more than 20 bp with Phred quality scores $<20$, homopolymer runs longer than 30 bp, or at least one 12-mer match to Illumina adaptor sequences. Reads were trimmed to remove non-template bases introduced during library prep, identified based on occurrence of a 'GGG' motif within the first 17 bp. Reads were then mapped to the set of 34,211 primary transcripts from the *S. bicolor* v3.1 genome with SHRiMP v.2.2.3 (32) and we kept only unique alignments and those with more than 40 bp matching the reference.

<u>Oligos used in TagSeq library preparations:</u>
**TruSeq_Un1:** AAT GAT ACG GCG ACC ACC GAG ATC TAC ACA TCA CGA CAC TCT TTC CCT ACA CGA CGC TCT TCC GAT CT
**TruSeq_Un2:** AAT GAT ACG GCG ACC ACC GAG ATC TAC ACA CTT GAA CAC TCT TTC CCT ACA CGA CGC TCT TCC GAT CT
**ILL-BC** (8 six-mer barcodes, here shown as NNNNNN):
CAAGCAGAAGACGGCATACGAGATNNNNNNGTGACTGGAGTTCAGACGTGTGCTCT TCCGATC
**S-Ill-swMW:**
rArCrCrCrCrArUrGrGrGrGrCrUrArCrArCrGrArCrGrCrUrCrUrUrCrCrGrArUrCrUrNrNMWr GrGrG
**3ILL-30TV:** ACGTGTGCTCTTCCGATCTAATTTTTTTTTTTTTTTTTTTTTTTTTTTTTTV
**5ILL:** CTACACGACGCTCTTCCGATCT

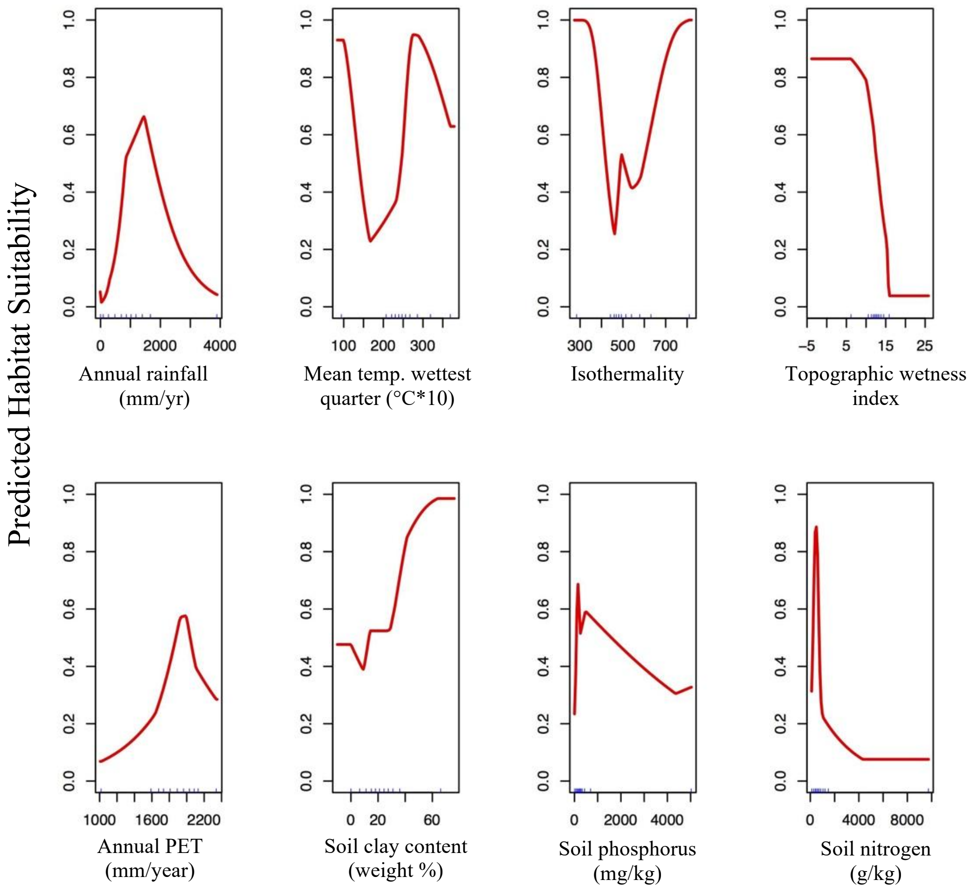


**Fig. S1.** Response curves for 8 environmental variables in all occurrence *S. hermonthica* distribution model. Curves represent the change in predicted habitat suitability (y-axis) for each variable with all other variables held at a constant value.

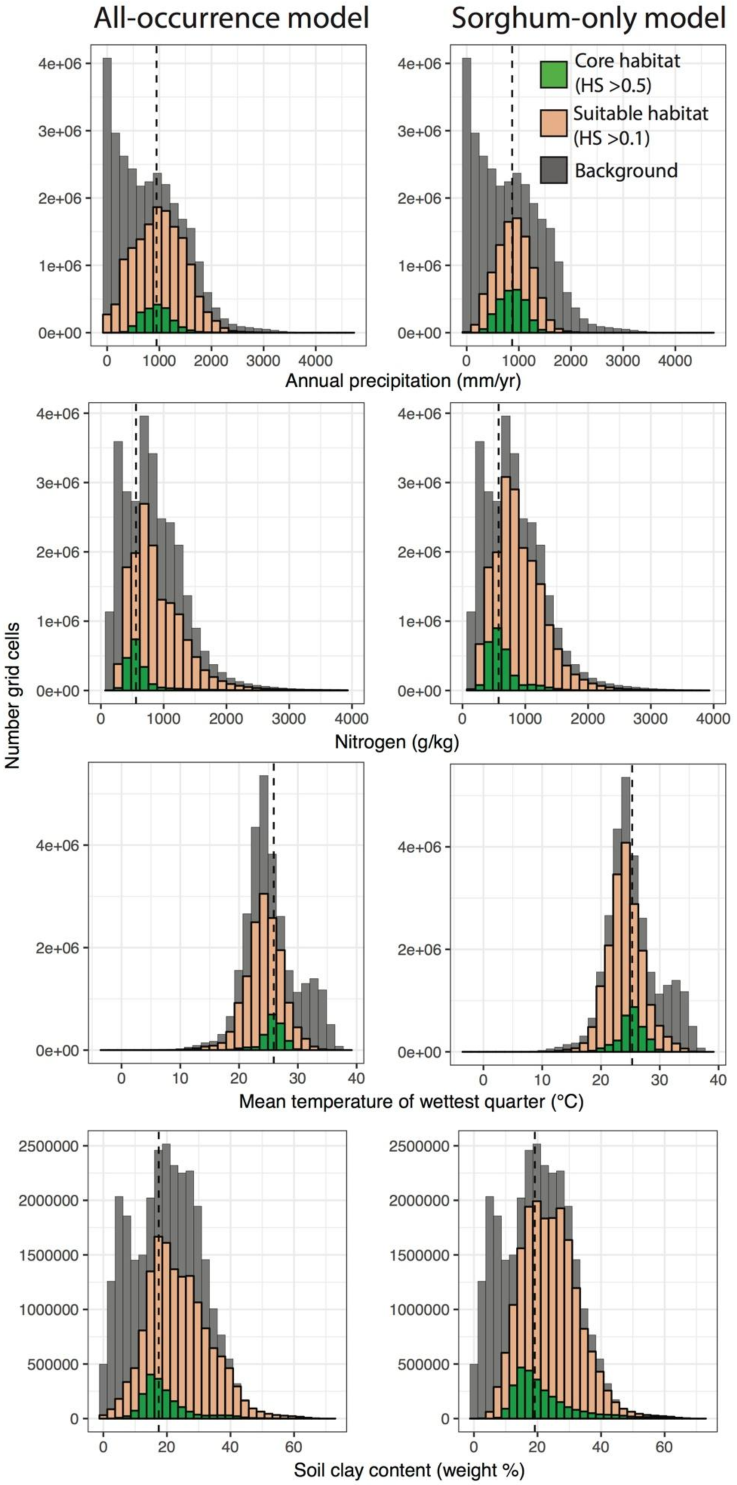


**Fig. S2.** Distribution of environmental values for four variables with permutation importance greater than 10% for the all-occurrence or sorghum-only *S. hermonthica* SDMs. Distributions are shown for all grids cells exceeding habitat suitability (HS) scores of 0.5 or 0.1, or for all grid cells in the background.

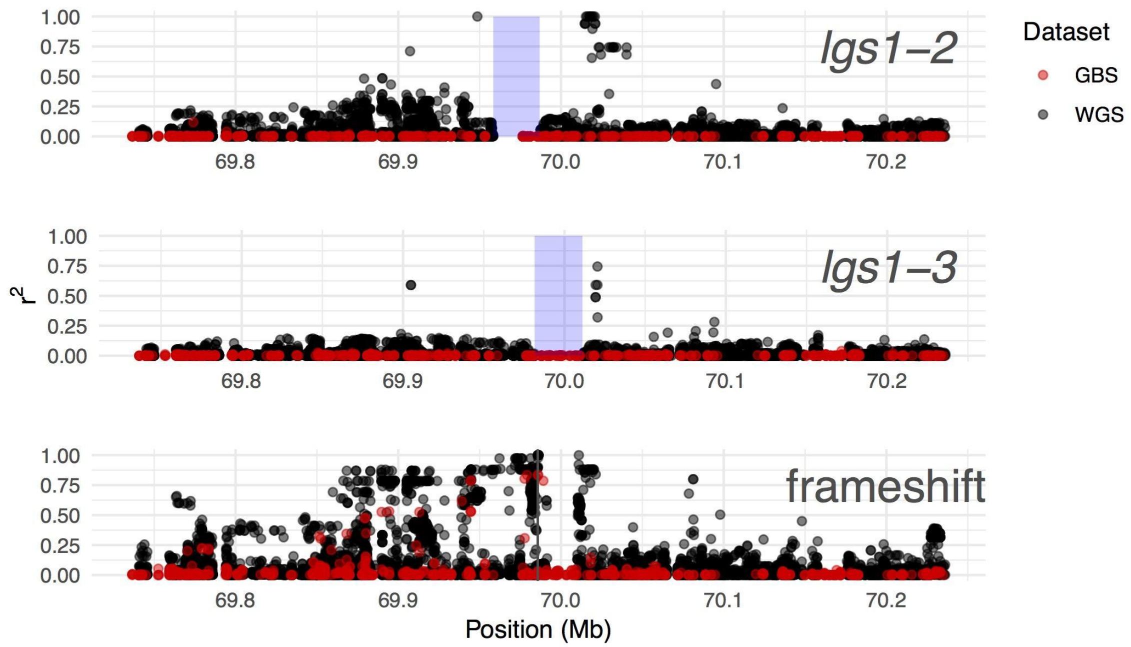


**Fig. S3.** Linkage disequilibrium with large deletion variants and the frameshift mutation at position 69,986,146 on *S. bicolor* Chromosome 5. Shaded boxes indicate positions of deletions (*lgs1-2*: 69,958,377-69,986,892*; lgs1-3*: 69,981,502-70,011,149) and vertical line indicates position of SNP tagging the frameshift mutation in the GBS dataset. For the WGS dataset, we excluded genotypes with depth of coverage less than 5x and SNPs with >70% missing data. Due to large amounts of missing data in the GBS dataset, $r^2$ was calculated based on imputed SNPs. An 18.8kb deletion beginning at position 69,991,579 also occurs in accessions with the frameshift, leading to missing data in this region for the non-imputed (WGS) dataset.

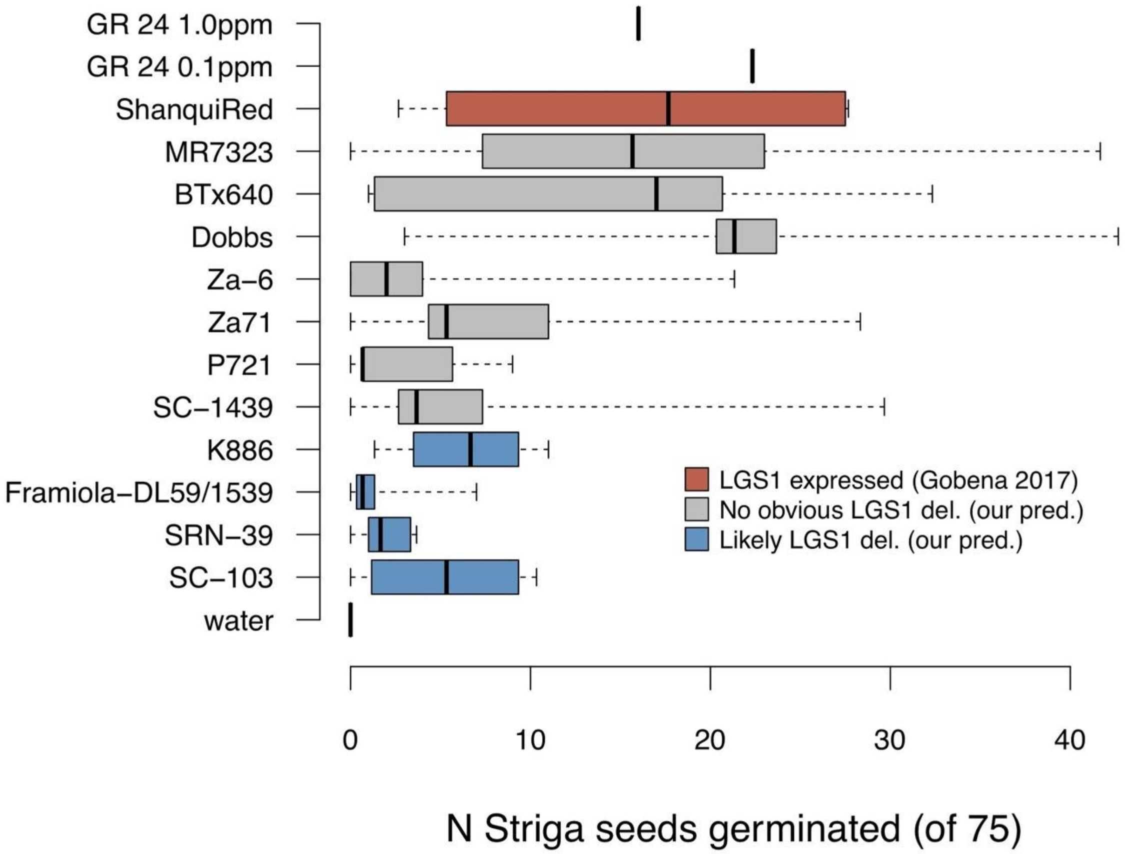


**Fig. S4.** Germination of *S. hermonthica* in response to root exudates isolated from 12 sorghum varieties. Boxes show interquartile range and median, whiskers show maximum and minimum. Predictions for LGS1 deletions are based on genotypes from the GBS dataset.

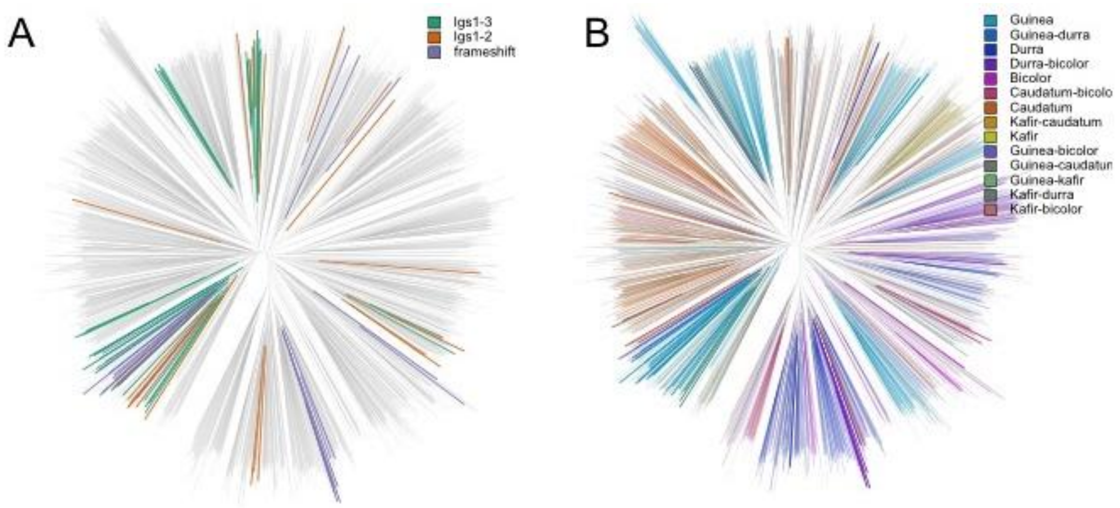


**Fig. S5.** Unrooted neighbor joining tree for *S. bicolor* chromosome 5 based on 2070 georeferenced individuals and 5434 SNPs with MAF >1%, after removing SNPs with >30% missing genotype calls. Branches are color-coded according to LGS1 deletion allele (A) or botanical race (B).

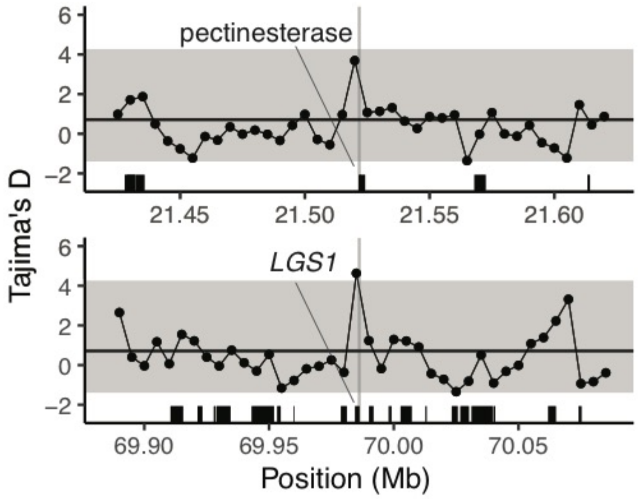


**Fig. S6. Signatures of balancing selection surround parasite-associated SNPs in a pectinesterase gene and *LGS1*.** Tajima's D was calculated in non-overlapping windows of 5 kb using data from the WGS dataset subset to 143 African landraces. The horizontal line indicates the median value for 1000 randomly selected 5 kb windows overlapping or encompassing gene models, and values overlapping the central 95% of the distribution are shown in the grey shaded region. Vertical lines indicate positions of parasite-associated SNPs on *S. bicolor* Chromosomes 2 (S2_21521798) and 5 (S5_69986146).

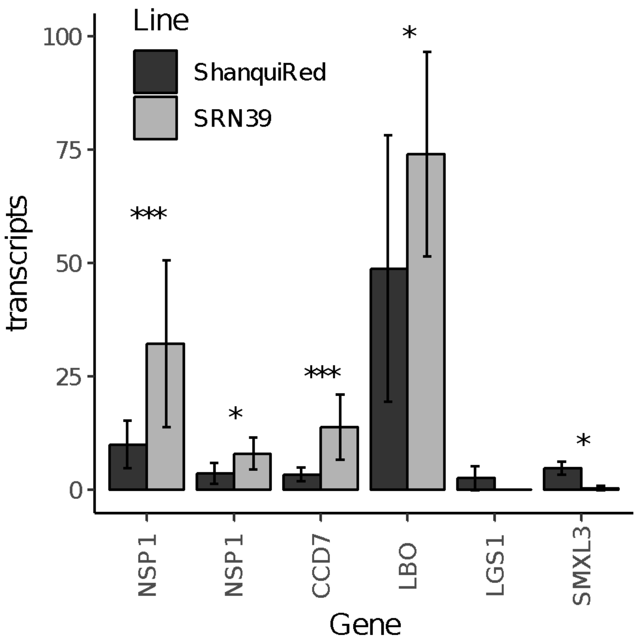


**Fig. S7**. Strigolactone biosynthesis and signaling pathway genes differentially regulated between nutrient-stressed roots of *LGS1*-deficient sorghum line SRN39 and *LGS1*-intact line Shanqui Red. Mean and standard deviation of read counts from 5 biological replicates per line are shown. Asterisks indicate significance following FDR correction. See Table S3 for corresponding gene models. ***$p < 0.001$; **$p < 0.01$; *$p < 0.05$.

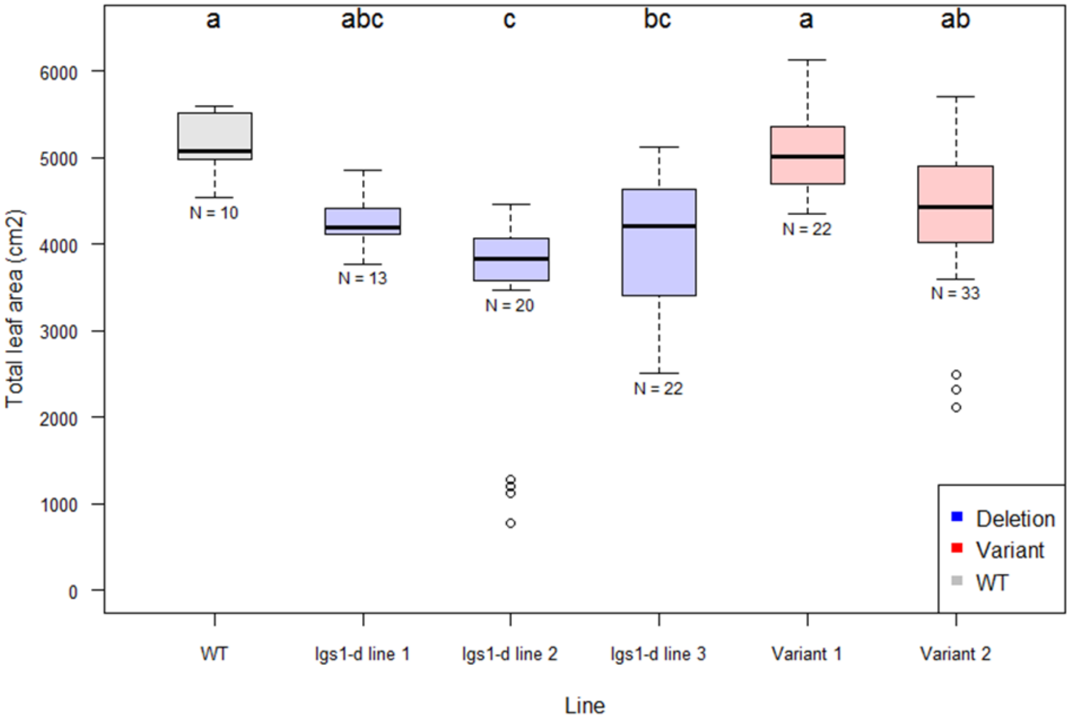


**Fig. S8**. Total leaf area at 65 days after planting. WT, wild-type Macia control. The CRISPR-cas *lgs1* deletion lines originated from three independent T0 plants. The *lgs1-d* line 3 and Variant 1 were derived from the same T0 plant. Variant 1 and Variant 2 contain intact *LGS1*, but have SNPs at the two cutting sites flanking the gene. Results of a one-way ANOVA followed by Scheffé's method are indicated by the letters above the box plots, where different letters indicate significant differences ($p < 0.05$). Sample size (N) is indicated below each box plot.

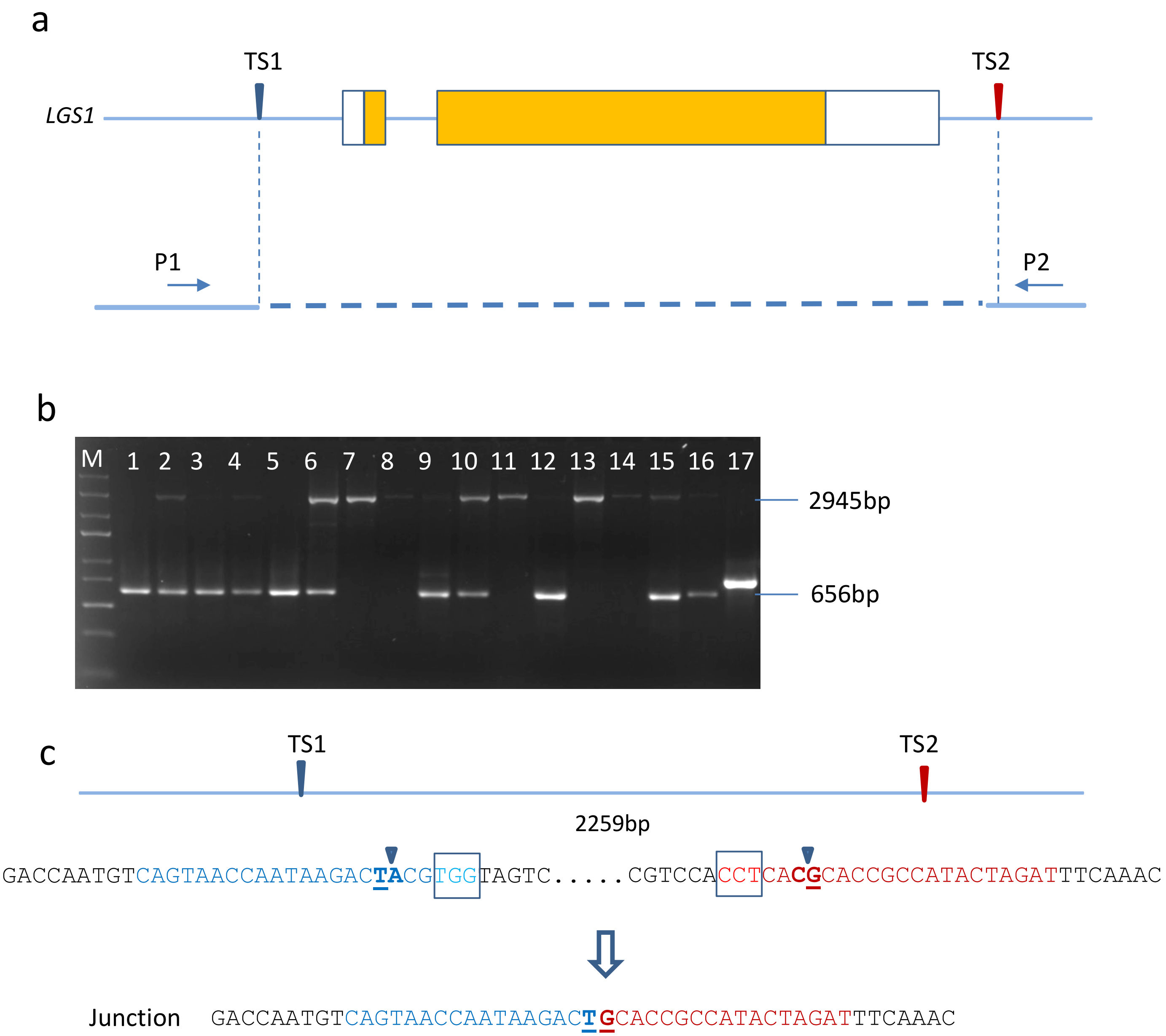


**Figure S9. *LGS1* gene deletion in Macia**

a) Schematic illustration of the sorghum *LGS1* gene. TS, CRISPR target site. The relative position and direction of PCR primers are indicated by arrows labelled P1 and P2. (b) Example gel image for deletion screening. Lane M, GeneRuler Express DNA Ladder. (c) Junction sequence of the *LGS1* deletion lines. Blue and red fonts indicate the TS1 and TS2 target sequences, respectively. PAM sequence is boxed. Arrows indicate the predicted Cas9 cleavage positions. Underlined nucleotides indicate the position where precise end-joining occurred at predicted cleavage sites.

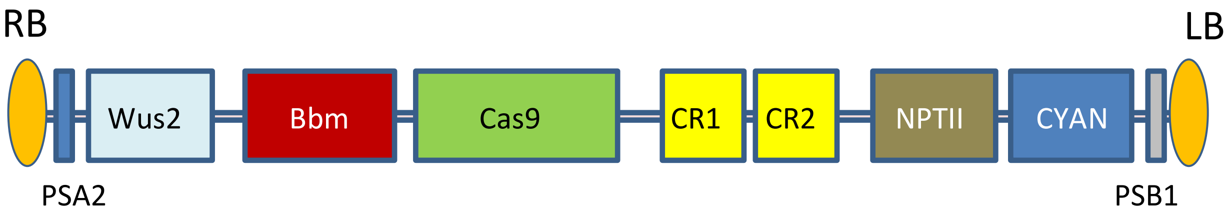


**Figure S10. Schematic drawing illustrating T-DNA region.**
WUS2 was under control of the maize Axig promoter. The expression of BBM was driven by maize Pltp promoter. Cas9 and NPTII were driven by the maize UBI promoter, guide RNA CR1 and CR2 under the maize U6 promoter, and Cyan under the Ltp promoter. PSA1 and PSB1, unique sequences to facilitate event screening.

**Table S1. Environmental predictors for species distribution models.** Core ranges are given as the 10th and 90th percentiles for grid cell values within the study extent (18W to 62E; 37S to 38N) with habitat suitability score >0.5 (all-occurrence and sorghum-only models) or for all grid cells within the full study extent. Median values are shown in parentheses. Models showed good discrimination ability with AUC values of 0.86 (all-occurrence model) or 0.85 (sorghum-only model). Both models included linear, quadratic, hinge, and product feature classes with a regularization multiplier of 1 for the all-occurrence model or 4 for the sorghum-only model. PI: Permutation importance.

| Variable | Units | Source | PI, % (all-occurrence) | PI, % (sorghum-only) | Core range (all-occurrence | Core range (sorghum-only) | Core range (full extent) |
|---|---|---|---|---|---|---|---|
| Annual rainfall (Bio12) | mm/yr | CHELSA | 35.6 | 28.7 | 608-1307 (947) | 543-1183 (871) | 42-1665 (737) |
| Total nitrogen at 0-30 cm depth (NTO) | g/kg | AfSoilGrids250m | 24.3 | 28.8 | 411-932 (559) | 399-977 (575) | 257-1449 (766) |
| Clay fraction at 5 cm depth (CLYPPT) | weight % | SoilGrids250m | 4.6 | 24.0 | 12-29 (17) | 13-36 (19) | 5-34 (20) |
| Mean temperature of wettest quarter (Bio8) | °C | CHELSA | 15.4 | 4.0 | 23.8-27.8 (25.9) | 22.5-27.8 (25.3) | 20.6-32.3 (24.7) |
| Isothermality (Bio3) | - | CHELSA | 8.1 | 4.8 | 464-583 (485) | 464-573 (486) | 433-622 (491) |
| Topographic wetness index (topoWet) | - | ENVIREM | 6.5 | 6.6 | 11.1-13.5 (12.5) | 10.5-13.8 (12.5) | 10.4-14.6 (12.6) |
| Potential evapotranspiration (annualPET) | mm/yr | ENVIREM | 4.0 | 2.9 | 1774-2044 (1937) | 1789-2090 (1958) | 1540-2126 (1821) |
| Total phosphorus at 0-30 cm depth | mg/kg | AfSoilGrids250m | 1.6 | 0.2 | 117-426 (192) | 119-465 (211) | 63-653 (264) |

**Table S2.** ***LGS1* loss-of-function mutations for African landraces in the WGS dataset.** Start positions are for Chromosome 5 of the *Sorghum bicolor* v3.0 genome.

| Accession | Origin | Race | Variant type | Variant size (bp) | Start | Genotype |
|---|---|---|---|---|---|---|
| PI221651 | Nigeria | Guinea | Insertion | 2 | 69986146 | 1/1 |
| PI221651 | Nigeria | Guinea | Deletion | 315 | 69984268 | 1/1 |
| PI329338 | Ethiopia | Caudatum-bicolor | Insertion | 2 | 69986146 | 1/1 |
| PI562781 | Mali | Guinea | Deletion | 29647 | 69981502 | 1/1 |
| PI562971 | Nigeria | Other | Deletion | 28515 | 69958377 | 1/1 |
| PI562981 | Nigeria | Other | Deletion | 28515 | 69958377 | 1/1 |
| PI562982 | Nigeria | Guinea | Deletion | 28515 | 69958377 | 1/1 |
| PI562990 | Nigeria | Other | Insertion | 2 | 69986146 | 1/1 |
| PI562990 | Nigeria | Other | Deletion | 315 | 69984268 | 1/1 |
| PI562991 | Nigeria | Other | Deletion | 28515 | 69958377 | 1/1 |
| PI562994 | Nigeria | Other | Deletion | 28515 | 69958377 | 1/1 |
| PI562998 | Nigeria | Other | Insertion | 2 | 69986146 | 1/1 |
| PI563002 | Nigeria | Other | Insertion | 2 | 69986146 | 1/1 |
| PI563009 | Nigeria | Guinea-caudatum | Deletion | 29647 | 69981502 | 1/1 |
| PI563020 | Nigeria | Other | Deletion | 29647 | 69981502 | 1/1 |
| PI563021 | Nigeria | Unknown | Insertion | 2 | 69986146 | 1/1 |
| PI563021 | Nigeria | Unknown | Deletion | 315 | 69984268 | 0/1 |
| PI563022 | Nigeria | Other | Insertion | 2 | 69986146 | 1/1 |
| PI563022 | Nigeria | Other | Deletion | 315 | 69984268 | 1/1 |
| PI585406 | Nigeria | Guinea | Insertion | 2 | 69986146 | 1/1 |
| PI585406 | Nigeria | Guinea | Deletion | 315 | 69984268 | 1/1 |
| PI585448 | Ghana | Guinea | Insertion | 2 | 69986146 | 1/1 |
| PI585448 | Ghana | Guinea | Deletion | 315 | 69984268 | 1/1 |
| PI585448 | Ghana | Guinea | Deletion | 29647 | 69981502 | 0/1 |
| PI585452 | Ghana | Guinea | Insertion | 2 | 69986146 | 0/1 |
| PI585452 | Ghana | Guinea | Deletion | 315 | 69984268 | 0/1 |
| PI585467 | Ghana | Guinea | Insertion | 2 | 69986146 | 1/1 |
| PI585467 | Ghana | Guinea | Deletion | 315 | 69984268 | 1/1 |
| PI585966 | Togo | Guinea | Insertion | 2 | 69986146 | 1/1 |
| PI585966 | Togo | Guinea | Deletion | 315 | 69984268 | 1/1 |

**Table S3. Sorghum genes related to strigolactone biosynthesis or signalling.** The number of high impact variants with minor allele frequency (MAF) >5% in the WGS dataset are shown. Gene models with less than 5 SNPs were excluded from Tajima's D calculations. Expression for each gene model in root tissue of sorghum lines Shanqui Red and SRN39 are shown as the mean count for reads mapped, with asterisks indicating significant differences between lines. **adjusted $p$ <0.001; *adjusted $p$ <0.05

| Homolog | Gene model | Function | Chr. | Pos. | Tajima's D | High impact variants (MAF >5%) | Expression |
|---|---|---|---|---|---|---|---|
| *NSP1* | Sobic.001 G341400 | GRAS-family transcription factor | 1 | 62,864,836-62,867,833 | NA | 0 | 21.3** |
| *NSP2* | Sobic.001 G428600 | GRAS-family transcription factor | 1 | 70,751,978-70,753,672 | -0.51 | 0 | 3.0 |
| *D14* | Sobic.001 G465100 | α/β-hydrolase | 1 | 73,888,745-73,895,024 | 0.96 | 0 | 253 |
| *NSP1* | Sobic.002 G372100 | GRAS-family transcription factor | 2 | 73,049,658-73,053,841 | -0.90 | 0 | 5.9* |
| *CCD8* | Sobic.002 G380550 | carlactone synthase | 2 | 73,655,874-73,656,496 | -0.84 | 1 | 0 |
| *MAX1* | Sobic.003 G269500 | cytochrome P450 | 3 | 60,624,004-60,628,867 | 1.28 | 0 | 38.2 |
| *MAX1* | Sobic.003 G269600 | cytochrome P450 | 3 | 60,633,944-60,636,136 | 0.92 | 1 | 73.4 |
| *CCD8* | Sobic.003 G293600 | carlactone synthase | 3 | 62,602,964-62,607,023 | -0.23 | 0 | 24.0 |
| *LBO* | Sobic.003 G418000 | oxidoreducta se | 3 | 72,382,455-72,384,461 | 3.80 | 0 | 60.6* |
| *MAX1* | Sobic.004 G095500 | cytochrome P450 | 4 | 8,304,449-8,307,025 | 2.39 | 0 | 0.4 |
| *SMXL5* | Sobic.004 G139900 | Class I Clp ATPase | 4 | 40,568,128-40,571,634 | -0.07 | 0 | 1.5 |
| *SMXL3* | Sobic.004 G168600 | Class I Clp ATPase | 4 | 51,922,148-51,925,516 | -0.19 | 2 | 1.9 |
| *SMAX1* | Sobic.004 G325000 | Class I Clp ATPase | 4 | 66,043,348-66,044,531 | 0.54 | 0 | 57.0 |

| | | | | | | | |
|---|---|---|---|---|---|---|---|
| *SMXL7* | Sobic.005 G002400 | Class I Clp ATPase | 5 | 190,488-194,282 | -0.64 | 0 | 3.9 |
| *CCD8* | Sobic.005 G002500 | carlactone synthase | 5 | 202,329-204,425 | -1.41 | 4 | 0.26 |
| *SMXL3* | Sobic.005 G043000 | Class I Clp ATPase | 5 | 4,061,603-4,065,225[1] | 0.74 | 3 | 24.3 |
| *CCD8* | Sobic.005 G105700 | carlactone synthase | 5 | 19,797,305-19,799,526 | -2.15 | 0 | 1.4 |
| *D27* | Sobic.005 G168200 | β-carotene isomerase | 5 | 64,672,109-64,677,619 | -0.88 | 0 | 3.0 |
| *LGS1* | Sobic.005 G213600 | sulfotransfera se | 5 | 69,984,440-69,986,256 | 0.19 | 3 | 1.1 |
| *SMXL3* | Sobic.006 G074900 | Class I Clp ATPase | 6 | 43,746,994-43,751,171 | -0.13 | 0 | 4.5 |
| *CCD7* | Sobic.006 G170300 | β-carotene cleaving dioxygenase | 6 | 52,713,305-52,716,037[2] | 0.47 | 2 | 9.6** |
| *D27* | Sobic.007 G016600 | β-carotene isomerase | 7 | 1,433,383-1,435,073 | -1.13 | 0 | 0 |
| *SMAX1* | Sobic.007 G090900 | Class I Clp ATPase | 7 | 14,458,102-14,461,202 | 2.31 | 0 | 80.6 |
| *CCD8* | Sobic.007 G170300 | carlactone synthase | 7 | 60,503,029-60,506,362 | -1.03 | 0 | 17.2 |
| *SMXL7* | Sobic.008 G002400 | Class I Clp ATPase | 8 | 219,391-225,260 | 0.26 | 0 | 245.2 |
| *SMXL3* | Sobic.008 G042800 | Class I Clp ATPase | 8 | 4,207,967-4,210,364 | NA | 0 | 2.2* |
| *NSP2* | Sobic.008 G046700 | GRAS-family transcription factor | 8 | 4,591,222-4,596,201 | -1.58 | 0 | 0.4 |
| *D27* | Sobic.009 G030800 | β-carotene isomerase | 9 | 2,741,604-2,744,934 | NA | 0 | 13.9 |

[1] Overlaps with $QTL05_1$ (11)

[2] Overlaps with QTL06 (11)

| | | | | | | | |
|---|---|---|---|---|---|---|---|
| *MAX2* | Sobic.010 G043000 | F-box protein | 10 | 3,312,894-3,316,043 | NA | 0 | 222.1 |
| *MAX1* | Sobic.010 G170400 | cytochrome P450 | 10 | 50,177,361-50,180,730 | NA | 0 | 33.7 |

*D27: DWARF27; CCD7: CAROTENOID CLEAVING DIOXYGENASE 7; CCD8: CAROTENOID CLEAVING DIOXYGENASE 8; MAX1: MORE AXILLARY BRANCHES 1; NSP1: NODULATION SIGNALING PATHWAY 1; NSP2: NODULATION SIGNALING PATHWAY 2; LBO: LATERAL BRANCHING OXIDOREDUCTASE; LGS1: LOW GERMINATION STIMULANT 1; SMAX1: SUPPRESSOR OF MAX2-1; SMXL7: SMAX1-LIKE 7; SMXL5: SMAX1-LIKE 5; SMXL3: SMAX1-LIKE 3*

**Table S4. Sorghum SNPs showing significant genome-wide associations with *S. hermonthica* distribution (FDR correction at $\alpha$=0.05).** SNPs within 1kb of gene models are annotated and are ordered according to genomic location. AF: reference allele frequency; AdjP: P-value after FDR adjustment. For gene models with more than one SNP, the highest scoring position is shown. SNPs predicted to result in an amino acid change are shown in bold.

| Chr | Position | AF | AdjP | Gene_model | Annotation |
|---|---|---|---|---|---|
| 1 | 21828170[3] | 0.292 | 0.046 | Sobic.001G227800 | Uncharacterized conserved protein, putative, expressed |
| 1 | 50469815 | 0.035 | 0.037 | NA | NA |
| 1 | 54864458[4] | 0.013 | 0.024 | NA | NA |
| 1 | 56800905 | 0.043 | 0.028 | NA | NA |
| 1 | 58808651 | 0.214 | 0.046 | NA | NA |
| 1 | 58837130 | 0.14 | 0.045 | Sobic.001G304500 | Os10g0551200 protein; GRAS domain family |
| 1 | 64142276 | 0.052 | 0.024 | NA | NA |
| 1 | 64164495 | 0.054 | 0.004 | Sobic.001G352600 | Expressed protein |
| 1 | 71929093 | 0.036 | 0.024 | Sobic.001G441500 | NA |
| 2 | 12723171 | 0.028 | 0.035 | NA | NA |
| 2 | 21521798 | 0.275 | 0.013 | Sobic.002G138400 | Pectinesterase 14-related |
| 2 | 68524689 | 0.018 | 0.049 | Sobic.002G311400 | Putative glucosyltransferase |
| 3 | 1842885 | 0.011 | 0.045 | NA | NA |
| 3 | 2025341 | 0.012 | 0.002 | Sobic.003G023500 | Os06g0116300 protein; putative syntaxin |
| 3 | 13445493 | 0.026 | 0.041 | Sobic.003G139000 | Os01g0290600 protein; putative L-cysteine desulfhydrase 2 |
| 3 | 51292869 | 0.012 | 0.023 | NA | NA |
| 3 | 56946323 | 0.055 | 0.043 | Sobic.003G229700 | glycerol-3-phosphate acyltransferase |
| 3 | 59075308 | 0.018 | 0.008 | Sobic.003G252200 | HB1, ASXL, restriction endonuclease HTH domain (HARE-HTH) |
| 3 | 62473154 | 0.077 | 0.002 | NA | NA |
| 3 | 62493603 | 0.093 | 0.028 | Sobic.003G292500 | ATP-dependent protease La (LON) domain-containing protein-like |
| 3 | 62795897 | 0.012 | 0.002 | Sobic.003G295900 | Putative heat shock factor |
| 3 | 67782723 | 0.012 | 0.008 | Sobic.003G360200 | WUSCHEL-related homeobox 9 |
| 3 | 70103330 | 0.055 | 0.017 | Sobic.003G389700 | Os01g0898800 protein |
| 4 | 19466351 | 0.011 | 0.045 | NA | NA |
| 4 | 19724561 | 0.011 | 0.028 | NA | NA |

[3] QTL from N13/E36-1 (33)

[4] QTL from N13/E36-1 (33)

| | | | | | |
|---|---|---|---|---|---|
| 4 | 19735015 | 0.011 | 0.020 | Sobic.004G132200 | NA |
| 4 | 20033536 | 0.014 | 0.040 | NA | NA |
| 4 | 20035072 | 0.012 | 0.027 | NA | NA |
| 4 | 20782140 | 0.012 | 0.012 | NA | NA |
| 4 | 23221234 | 0.041 | 0.012 | NA | NA |
| 4 | 60099891 | 0.012 | 0.002 | Sobic.004G254500 | Leafy cotyledon1 |
| 4 | 60767527 | 0.012 | 0.015 | NA | NA |
| 4 | 66105213 | 0.013 | 0.006 | Sobic.004G325832 | NA |
| 4 | 66766354 | 0.03 | 0.040 | NA | NA |
| 5 | 1950501 | 0.087 | 0.045 | NA | NA |
| 5 | 1976589 | 0.086 | 0.013 | Sobic.005G021400 | NA |
| 5 | 11089407 | 0.062 | 0.024 | NA | NA |
| 5 | 71122163 | 0.011 | 0.012 | Sobic.005G224300 | similar to O-methyltransferase |
| 6 | 487441 | 0.086 | 0.040 | Sobic.006G002900 | similar to OSJNBa0094O15.15 protein; zinc/RING finger domain |
| 6 | 15445897 | 0.011 | 0.046 | NA | NA |
| 6 | 42246344 | 0.129 | 0.028 | NA | NA |
| 6 | 43653802 | 0.022 | 0.018 | Sobic.006G074300 | similar to OSIGBa0092M08.2 protein; non-specific lipid transfer protein d6 |
| 6 | 51034850 | 0.028 | 0.033 | NA | NA |
| 6 | 51042152 | 0.079 | 0.050 | Sobic.006G148800 | Phenylalanine ammonia-lyase |
| 6 | 52891272 | 0.087 | 0.004 | NA | NA |
| 6 | 59662162 | 0.012 | 0.042 | Sobic.006G261700 | similar to OSJNBb0004A17.4; Lactoylglutathione lyase |
| 7 | 11575066 | 0.014 | 0.018 | NA | NA |
| 7 | 12044293 | 0.014 | 0.016 | NA | NA |
| 7 | 12094853 | 0.013 | 0.012 | NA | NA |
| 7 | 12891407 | 0.019 | 0.045 | NA | NA |
| 7 | 12895894 | 0.01 | 0.015 | NA | NA |
| 7 | 14193502 | 0.018 | 0.045 | NA | NA |
| 7 | 14220105 | 0.018 | 0.034 | NA | NA |
| 7 | 14279216 | 0.015 | 0.028 | NA | NA |
| 7 | 14459084[5] | 0.014 | 0.007 | Sobic.007G090900 | ATP dependent CLP protease; SMAX1 |
| 7 | 14942059 | 0.018 | 0.026 | Sobic.007G091200 | similar to H0117D06-OSIGBa0088B06.9 protein; Long-chain-alcohol O-fatty- |

[5] SL signaling (SMAX1 ortholog)

| | | | | | |
|---|---|---|---|---|---|
| | | | | | acyltransferase |
| 7 | 15219432 | 0.021 | 0.040 | Sobic.007G091400 | diacylglycerol kinase (ATP) (dgkA, DGK) |
| 7 | 15254559 | 0.013 | 0.004 | NA | NA |
| 7 | 15325565 | 0.017 | 0.045 | NA | NA |
| 7 | 15359204 | 0.013 | 0.024 | NA | NA |
| 7 | 15532403 | 0.013 | 0.012 | NA | NA |
| 7 | 15534152 | 0.013 | 0.004 | NA | NA |
| 7 | 15554515 | 0.012 | 0.002 | Sobic.007G092000 | CCT/B-box zinc finger protein |
| 7 | 15653673 | 0.019 | 0.046 | Sobic.007G092200 | aspartate carbamoyltransferase |
| 7 | 15849206 | 0.014 | 0.027 | NA | NA |
| 7 | 15901805 | 0.017 | 0.026 | NA | NA |
| 7 | 15929687 | 0.019 | 0.018 | Sobic.007G092600 | Receptor-like kinase Xa21-binding protein 3-like |
| 7 | 16293907 | 0.014 | 0.027 | NA | NA |
| 7 | 17082700 | 0.014 | 0.044 | NA | NA |
| 7 | 17092629 | 0.017 | 0.040 | Sobic.007G093900 | N-terminal acetyltransferase |
| 7 | 17302379 | 0.019 | 0.045 | Sobic.007G094600 | Cytochrome P450 protein |
| 7 | 17831891 | 0.019 | 0.040 | Sobic.007G096200 | Putative uncharacterized protein |
| 7 | 17882047 | 0.018 | 0.028 | NA | NA |
| 7 | 17960172 | 0.018 | 0.018 | Sobic.007G096800 | Putative uncharacterized protein |
| 7 | 18275486 | 0.012 | 0.018 | NA | NA |
| 7 | 18279010 | 0.013 | 0.015 | NA | NA |
| 7 | 18300164 | 0.012 | 0.024 | NA | NA |
| 7 | 18408217 | 0.012 | 0.015 | NA | NA |
| 7 | 19041599 | 0.011 | 0.037 | Sobic.007G097400 | sin-like protein conserved region containing protein |
| 7 | 48195744 | 0.017 | 0.045 | NA | NA |
| 7 | 62605107 | 0.013 | 0.006 | Sobic.007G193500 | similar to Teosinte glume architecture 1 |
| 7 | 63109499 | 0.023 | 0.045 | NA | NA |
| 8 | 3177383 | 0.024 | 0.029 | Sobic.008G034400 | ATP-dependent RNA helicase, putative, expressed |
| 8 | 13233629 | 0.016 | 0.018 | NA | NA |
| 8 | 60143889 | 0.012 | 0.026 | Sobic.008G167300 | Leucine-rich repeat-containing protein |
| 8 | 61305329 | 0.014 | 0.043 | Sobic.008G179600 | similar to Expressed protein |
| 9 | 1792519 | 0.011 | 0.017 | Sobic.009G019900 | NA |
| 9 | 5744893 | 0.226 | 0.046 | Sobic.009G056600 | Putative uncharacterized protein |
| 9 | 49976645 | 0.014 | 0.028 | Sobic.009G142400 | Putative gibberellin 20-oxidase |

| | | | | | |
|---|---|---|---|---|---|
| 9 | 51474569 | 0.110 | 0.028 | NA | NA |
| 9 | 56421520 | 0.014 | 0.040 | NA | NA |
| 10 | 13677824 | 0.014 | 0.024 | NA | NA |
| 10 | 55681432 | 0.020 | 0.013 | Sobic.010G213900 | Receptor protein kinase PERK1-like |
| 10 | 56006575 | 0.014 | 0.014 | Sobic.010G216800 | Putative uncharacterized protein |
| 10 | 56761438[6] | 0.026 | 0.018 | NA | NA |
| 10 | 59272319 | 0.016 | 0.028 | Sobic.010G253700 | NA |

[6] QTL from IS9830/E36-1 (33)

**Table S5. Accessions from the Sorghum Association Panel used in germination study.**

| Accession | Name | Origin | Reported Resistance | *LGS1* allele |
|---|---|---|---|---|
| PI 656027 | SRN39 | - | Resistant | Deletion |
| PI 533752 | SC103 (54.K.94) | - | Resistant | Deletion |
| PI 533972 | Dobbs | Uganda | Resistant (post-attachment) | Intact |
| PI 656025 | Shanqui Red | China | Susceptible | Intact |
| PI 656055 | P721 | US | Susceptible | Intact* |
| PI 656051 | MR7323 | Niger | Unknown | Intact |
| PI 642791 | BTx640 | US | Unknown | Intact |
| PI 534088 | Za6 | Nigeria | Unknown | Intact |
| PI 534092 | Za71 | Nigeria | Unknown | Intact |
| PI 533976 | Framiola DL59/1539 | South Africa | Unknown | Deletion |
| PI 656040 | K886 | - | Unknown | Deletion |
| PI 656081 | SC1439 | - | Unknown | Intact* |

*No data in coding region, although SNP calls inconsistent with breakpoints for *lgs1-2* or *lgs1-3*.

**Table S6. Molecular characterization of S1 plants with *LGS1* deletion.**

| Line | Number of S1 plants | S1 Plant with *LGS1* deletion | *LGS1* deletion plant free of T-DNA | SbS |
|---|---|---|---|---|
| 1 | 32 | 25 | 3 | yes |
| 2 | 32 | 26 | 7 | yes |
| 3 | 32 | 25 | 5 | yes |

**Table S7. Molecular Inversion Probe (MIP) primers for Next-Generation Sequencing (NGS) the *LGS1* gene deletion junctions.**

| PCR | Primer | Sequence |
|---|---|---|
| 1st | P1f | ATCGGGAAGCTGAAGTTTCAAACAAAGTCCTCCTGCAATG |
| | p2r | ATCCGACGGTAGTGTTTGTTTAATTCTTTCATGTGGTTCTATTTGT |
| 2nd | 2pf | AATGATACGGCGACCACCGAGATCTACACATACGAGATCCGTAATCGGGAAGCTGAAG |
| | 2pr | CAAGCAGAAGACGGCATACGAGATNNNNNNNNACACGCACGATCCGACGGTAGTGT |

**Table S8. Primers and probes for detecting CRISPR-Cas9 reagents and helper genes.**

| Primer | Orientation | Sequence (5’ to 3’) |
|---|---|---|
| ADH-F | forward | CAAGTCGCGGTTTTCAATCA |
| ADH -R | reverse | TGAAGGTGGAAGTCCCAACAA |
| ADH- P | probe | VIC-TGGGAAGCCTATCTACCAC |
| Cas9-F | forward | CAGAATGAAAAGCTCTACCTCTACTACCT |
| Cas9-R | reverse | TGGTCGACGTCGTAGTCCGA |
| Cas9-P | probe | 6FAM-TCCTGGTCCACGTACAT |
| gRNA-F | forward | CTAATCACAAGAGTGGAGCGTACCTT |
| gRNA-R | reverse | AGCCTTATTTTAACTTGCTATTTCTAGCTCT |
| gRNA-P | probe | FAM-CCGAGCCGCAAGCA |
| NPTII-F | forward | CGTTGGCTACCCGTGATATTG |
| NPTII-R | reverse | GGAAGCGGTCAGCCCATT |
| NPTII-P | probe | FAM-TGAAGAGCTTGGCGGC |
| Bbm-F | forward | CGGCGATGTCTGCTTCAA |
| Bbm-R | reverse | AAGCTCTGATCCCCTCATGCT |
| Bbm-P | probe | FAM-ATCCCCCAAGATTG |
| Wus-F | forward | CCACGCACATGACCGCA |
| Wus-R | reverse | TGCCTCCTCCCGCTCC |
| Wus-P | probe | FAM-ACCGCCGCCCGCA |

**Additional data table S1 (separate file)**

CSV file of sorghum accessions from the GBS dataset with predicted *LGS1* loss-of-function alleles. Botanical race is coded as 'NA' if accession was not found in master list of accession annotations. Accessions with missing data at SNP S5_69985710 tagging the frameshift are not included in this list.

**Additional data table S2 (separate file)**

Excel file listing differentially expressed genes (FDR = 0.05) between nutrient-stressed roots of *Striga*-susceptible sorghum line Shanqui Red and *Striga*-resistant line SRN39 or between Macia wild-type and CRISPR *LGS1* knockout lines in roots or shoots. In addition, results for gene set enrichment analyses using ErmineJ for Macia vs. *LGS1* knockout lines are given for root and shoot tissues. Expression, fold change, and FDR corrected *p*-values are shown for all genes annotated with three GO terms that were significantly enriched among differentially expressed genes in roots or shoots and were discussed in the main text.

**Additional data table S3 (separate file)**

*Striga hermonthica* occurrence records used for species distribution models presented in this study.

**Additional data table S4 (separate file)**

Raster file of MaxEnt logistic output for all-occurrence *S. hermonthica* species distribution model.

**Additional data table S5 (separate file)**

Raster file of MaxEnt logistic output for sorghum-only *S. hermonthica* species distribution model.
